# Mitochondrial clone tracing within spatially intact human tissues

**DOI:** 10.1101/2025.07.11.664452

**Authors:** Sydney A. Bracht, Jiazhen Rong, Rodrigo A. Gier, Maureen DeMarshall, Hailey Golden, Diya Dhakal, Jayne C. McDevitt, Feiyan Mo, Emma E. Furth, Alexandra Strauss Starling, Amanda B. Muir, Gary W. Falk, Bryson W. Katona, Ben Z. Stanger, Nancy R. Zhang, Sydney M. Shaffer

## Abstract

Understanding tissue development and intra-tissue evolution requires the ability to trace clones in intact tissues coupled with high-plex molecular profiling preserving spatial context. However, current lineage tracing tools are incompatible with spatial omics. Here, we present SUMMIT (Spatially Unveiling Mitochondrial Mutations In Tissues), a spatially-resolved lineage tracing technology that integrates gene expression profiling with mitochondrial mutation-based clone identification. Unlike synthetic lineage recording methods, SUMMIT relies only on endogenous mutations and thus can be applied to human tissues. To address the compositional mixing of cell types within spatial spots, SUMMIT includes a rigorous statistical framework to confidently assign variants to specific cell subpopulations and achieves high power for spatially localized clones by pooling information across neighboring spots. We validated SUMMIT using a controlled model in which we mixed two cancer cell lines in a mouse tumor, then demonstrated it on multiple human tissues including Barrett’s esophagus, gastric cardia, small bowel, and colorectal cancer. Across these samples, we distinguished between global mutations and mutations marking locally restricted clones. The coupled transcriptomic data allowed us to characterize the cell type composition within each clone and delineate their spatial configuration. This integrated approach provides a framework to understand spatially-defined clonal evolution in preserved human tissue.

## Introduction

Understanding how tissues develop, maintain homeostasis, and undergo pathological transformation requires knowledge of both cellular lineage and the local microenvironmental interactions that shape cell behavior. Within a tissue, genetically distinct clones often coexist and follow divergent trajectories of growth and differentiation. This clonal diversification is especially consequential in cancer, where competition between clones drives disease progression and therapy resistance. Spatial transcriptomics technologies enable *in situ* mapping of gene expression and reveal cellular niches and tissue organization to study these processes; however, they provide only a static snapshot of cell state without lineage history. This is a major limitation, as a cell’s phenotype reflects both its ancestry and its current niche.

Recent developments in lineage tracing have enabled tracking of cellular ancestry alongside molecular features such as gene expression and chromatin accessibility. Synthetic lineage tracing relies on introducing DNA barcodes or CRISPR-editable arrays into the genome, which are passed down through cell divisions. While recent advances have enabled pairing of genetic barcoding with spatial transcriptomics or *in situ* sequencing^1,2,3^, these methods remain limited to *in vitro* systems or animal models. Alternatively, imaging-based lineage tracing preserves spatial contexts, but is restricted to short timescales and optically transparent specimens^4^. Retrospective lineage tracing techniques capture endogenous lineage specific features of cells derived from methylation^5,6^, mutations^7^, or copy number alterations^8,9^. While copy number alterations have recently been adapted for *in situ* lineage tracing using spatial transcriptomics^10,11^ and spatial genomics^12^ data, each represent only one modality of somatic change. The power of endogenous lineage tracing depends on the availability and density of naturally occurring lineage markers, and new methods are needed to uncover and leverage the full spectrum of somatic variation in intact tissues.

Mitochondrial DNA (mtDNA) offers an elegant solution to the challenges to endogenous lineage tracing. mtDNA mutations serve as ideal natural barcodes for reconstructing cellular relationships due to high copy number per cell, elevated mutation rate, and absence of recombination^13,14,15,16^. As cells divide, they inherit both nuclear and mitochondrial mutations, with the latter serving as stable markers of shared ancestry^17^. Mutations in mtDNA have been used to resolve lineages and identify stem cell niches using immunohistochemistry-based lineage tracing; however, this technique is limited to tracing mutations that affect cytochrome c oxidase (COX) expression^18,19^. More recently, single-cell mitochondrial genotyping methods have leveraged properties of mtDNA for lineage reconstruction using all mitochondrial mutations. However, these methods require tissue dissociation and thus sacrificing the spatial information critical for understanding tissue architecture and cell-cell interactions^20^.

To bridge this gap, we developed SUMMIT (Spatially Unveiling Mitochondrial Mutations In Tissue), a method that integrates mitochondrial lineage tracing with high resolution spatial transcriptomics. SUMMIT enables simultaneous visualization of gene expression patterns, cellular identities, and clonal relationships within their native tissue context by enriching mitochondrial transcripts from spatial cDNA libraries while preserving their spatial barcodes. To address the challenge of mixed cell types within spatial spots, SUMMIT incorporates a statistically rigorous framework to deconvolve lineage signals and assign variants to specific cell populations. We validate SUMMIT through controlled cell mixing experiments *in vivo* and demonstrate its utility across diverse human tissues, including Barrett’s esophagus, gastric cardia, small bowel, and colorectal cancer.

## Results

### Establishing proof of concept for spatial mitochondrial lineage tracing

Current mitochondrial lineage tracing techniques can be paired with single cell technologies, but cannot simultaneously read out molecular information and spatial location. To overcome this limitation, we developed SUMMIT to identify and spatially map clonal populations in intact human tissues. Here, we define a jclonej as a group of cells that share one or more identical mitochondrial mutations inherited from a common ancestral cell. These shared mutations act as natural barcodes that mark descendant cells of the same progenitor. Using these endogenous markers, SUMMIT addresses two central questions about tissue organization and function, 1) do clones segregate in space or intermingle, and 2) do cells of the same clone vary their transcriptional state with spatial context, and if so, how (Fig. 1A)? The SUMMIT workflow enables joint profiling of clonal mutations, transcriptional state, and spatial context from intact tissues (Fig. 1B).

**Figure 1:**
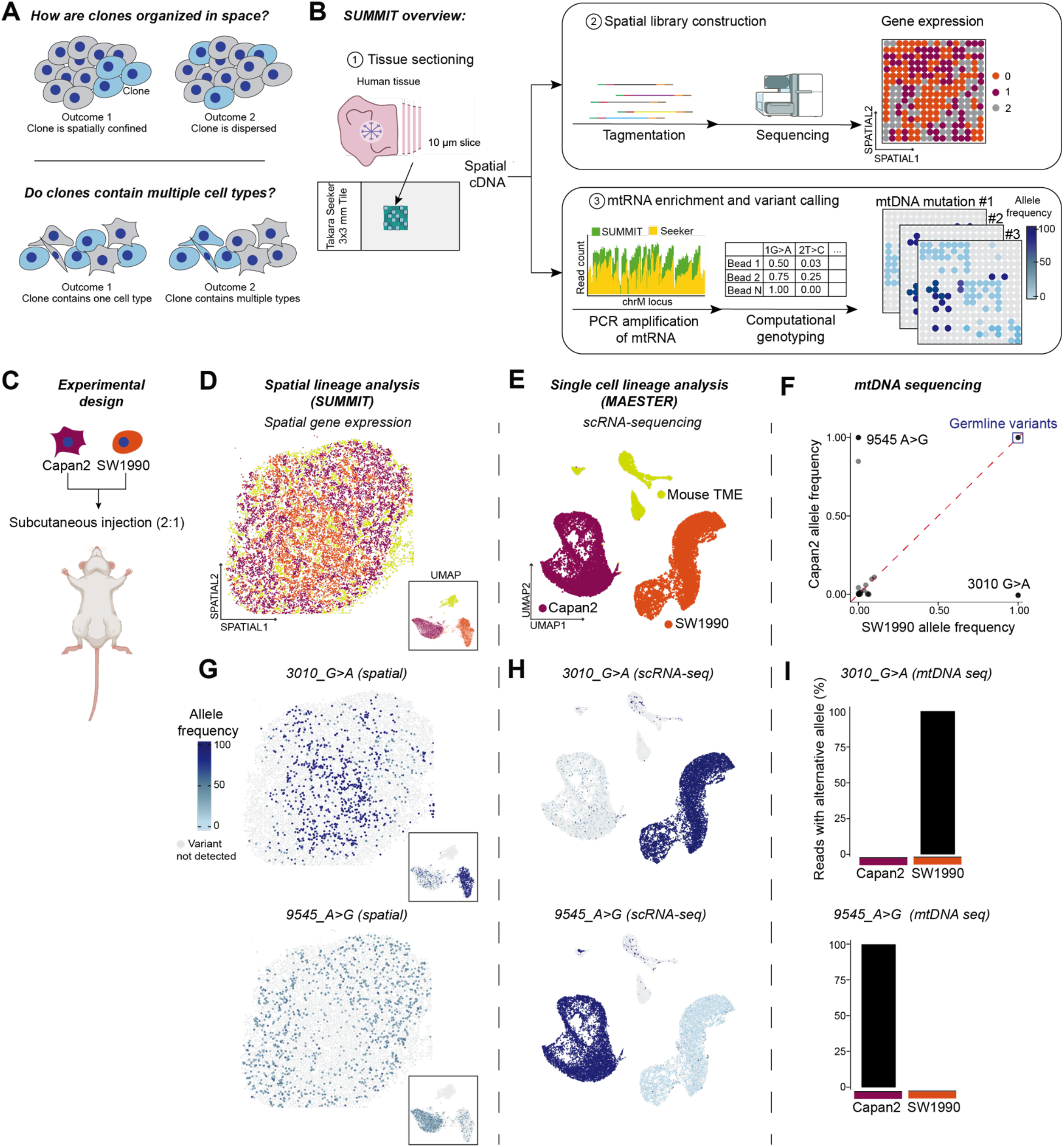
SUMMIT enables spatially resolved mitochondrial lineage tracing in intact tissues. (A) Conceptual motivation for spatial mitochondrial lineage tracing. B) SUMMIT workflow generates spatial cDNA for gene expression profiling and mitochondrial enrichment for variant calling, enabling joint analysis of transcriptional state and clonal relationships. (C) Experimental design using subcutaneous injection of human pancreatic cancer cell lines (Capan-2 and SW1990) in immunodeficient mice. (D) Spatial gene expression analysis showing segregation of Capan-2 and SW1990 determined by robust cell type deconvolution (RCTD). Single-cell RNA sequencing of parallel xenograft tumor showing distinct clustering of Capan-2, SW1990, and murine cells in UMAP space. (F) Mitochondrial DNA (mtDNA) sequencing identifies mutations distinguishing SW1990 and Capan-2. (G) Spatial mapping of mitochondrial variants 3010_G>A and 9545_A>G shows distinct spatial distributions corresponding to cell line locations. (H) Single-cell mitochondrial variant detection using MAESTER detects the same cell type specific variants. (I) Quantification of variant allele frequencies in mtDNA sequencing confirms cell line specificity.

To establish proof-of-concept for this technology, we designed a controlled experiment to test our ability to resolve known mitochondrial variants from a mixture of cell lines in intact tissues. We generated murine tumors by subcutaneously injecting a 2:1 mixture of two human pancreatic cancer cell lines (Capan-2 and SW1990) into immunodeficient mice (Fig. 1C). Seventeen days post-injection, we harvested the tumors and processed them in parallel through two workflows: (1) cryopreservation in OCT for spatial transcriptomics (Fig. 1D), and (2) immediate dissociation for single-cell RNA sequencing (scRNA-seq) (Fig. 1E). From an adjacent section, we collected DNA for mitochondrial DNA sequencing (Fig. 1F). This approach allowed us to directly compare the mitochondrial variants detected in spatial, single-cell, and DNA data using xenograft tumors from the same mouse.

scRNA-seq data confirmed that Capan-2 and SW1990 maintained distinct transcriptional profiles *in vivo*, appearing as well-separated clusters in UMAP visualization despite growing together in the same tumor (Fig. 1E). For spatial analysis, we used the Takara (formerly Curio) Seeker platform, which captures transcripts on 10 μm beads. While this resolution approaches single-cell scale, individual beads can capture RNA from multiple cells, particularly in densely packed tissues. Thus, to accurately estimate cell type composition within each bead, we performed robust cell type decomposition (RCTD)^21^. This computational deconvolution successfully resolved the tumor architecture, revealing spatial organization of the two human cancer cell lines with SW1990 forming an inner core surrounded by Capan-2 (Fig. 1D).

To enable mitochondrial mutation detection from spatial transcriptomics data, we adapted a targeted enrichment approach using PCR with primers tiling the mitochondrial genome, similar to the MAESTER method developed for scRNA-seq^20^. This enrichment step was critical because standard spatial transcriptomics libraries contain relatively few mitochondrial transcripts (mean coverage per locus = 1806.247, mean mitochondrial transcripts per cell = 2,248.513) and are insufficient for reliable variant calling. We systematically optimized three parameters to maximize mitochondrial transcript recovery: (1) initial cDNA amplification cycles, (2) cDNA input amount per PCR reaction, and (3) mitochondrial-specific PCR amplification cycles. Through this optimization, we determined that 6 initial PCR cycles, 10 ng cDNA input per reaction, and 6 mitochondrial-specific PCR cycles yielded optimal results (See Methods and Fig. S1A). The effectiveness of our enrichment strategy is demonstrated in Fig. S1B, where SUMMIT achieved 25-to 100-fold higher coverage per mitochondrial locus and transcripts per cell compared to the mitochondrial coverage in standard Takara Seeker libraries (mean reads per locus = 44,697.7, mean mitochondrial transcripts per cell = 55,642.08) without substantial PCR bias (Fig. S1C).

With this enhanced mitochondrial genome coverage, we performed variant calling on the SUMMIT-derived libraries and identified two variants, 3010_G>A and 9545_A>G, with distinct spatial distributions that corresponded precisely to the locations of the two human cell lines (Fig. 1G). The same variants were also detected in the parallel scRNA-seq data using MAESTER, confirming the consistency of our findings across library types (Fig. 1H). Additional cell line specific variants detected in mtDNA and scRNA-seq data (8002_C>T, 11152_T>C, 10176_G>A and 13500_T>C) were present in spatial data at insufficient coverage (Fig. S1D).

To validate that these variants are cell line specific, we performed mtDNA sequencing directly on each line and confirmed 3010_G>A to be specific to SW1990 and 9545_A>G to Capan-2^22^ (Fig. 1F, 1I). Together, these results demonstrate that SUMMIT can accurately capture and spatially resolve mitochondrial variants that mark distinct cellular populations from spatial transcriptomics libraries.

### Cell type-specific testing and spatial delineation of clones

A key challenge in analyzing spatial barcoding data is the potential mixing of cell types within spots^21,23,24^. Although the Takara Seeker platform offers 10m resolution, most spots contain mixtures of cells and subsequently, mtDNA variants (Fig. 2A, Fig. S1E). For instance, in the cell line mixing experiment, we observed that while the deconvolution algorithm could cleanly distinguish between Capan-2 and SW1990 cells (with only 1.3% mixed), both cell types frequently co-occurred with microenvironment cells in the same spots (22.1% for SW1990 and 23.2% for Capan-2) (Fig. S2A). To confidently assign mitochondrial mutations to their carrier cell types, we developed a computational pipeline within the SUMMIT framework (Fig. 2B, modules 1-3).

**Figure 2:**
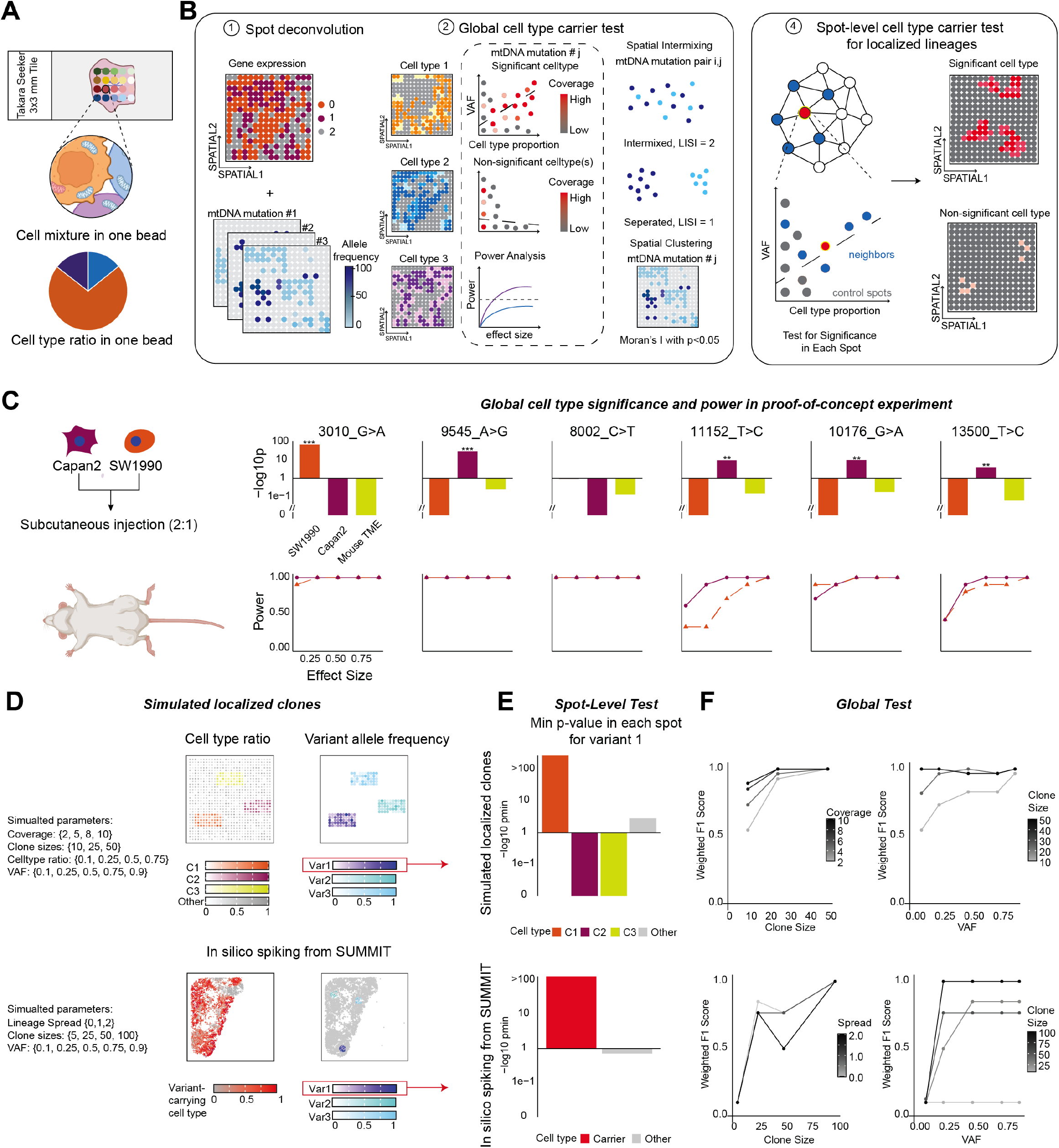
Statistical framework for cell type-specific variant assignment. (A) Challenge of cell type mixtures within one spatial spot. (B) SUMMIT computational pipeline workflow. Module 1: *Spot deconvolution using robust cell type deconvolution (RCTD) to estimate cell type pr*oportions. Module 2: Global cell type carrier test using regression of observed variant allele frequency against cell type proportions. Power analysis is paired with non-significant cell types to exclude the non-significant associations due to low coverage. Module 3: Spot-level significance testing with spatial smoothing for localized clone detection. Each spot is tested with neighboring spots and a set of control spots. (C) Validation of computational pipeline in xenograft experiment. Bar plots (top) show p-value from global significance testing results for each variant across cell lines. Variant 3010_G>A is significantly associated with SW1990, while variants 9545_A>G, 11152_T>C, 10176_G>A, and 13500_T>C are associated with Capan-2. Corresponding power analysis (bottom) demonstrates adequate statistical power for confident negative results. (D) Simulation framework design. (Top) Simulated localized clones with parameter changes in coverage, clone sizes, cell type ratio and VAF. (Bottom) In silico spiking from real SUMMIT data and biological structure with change in lineage spread, clone sizes and VAF. (E) Bar plots of minimum p-values for variant 1 (correspond to cell type 1) across cell types correctly identifying cell type 1 as the carrier. (F) F1 score of the pipeline performance across varying parameters in the simulations.

The pipeline takes as Input the estimated cell type proportions for each spot determined by RCTD^21^, mtDNA alternative allele counts, and total mtRNA detected per cell (Fig 2B, module 1). For each cell type and each mitochondrial variant, we perform a statistical test of global co-enrichment. Co-enrichment indicates the given cell type contains a nonzero fraction of variant-carrying cells. The test is based on a bead-level model, where the bead-level alternative allele frequency, conditioned on the total coverage, follows a mixture of Binomial distributions with success probability depending on the estimated spot-level contributions of the cell type. Operationally, this corresponds to a regression of the observed allele frequency against the deconvolved contribution of the cell type (Fig. 2B, module 2). Under the null hypothesis where no cells of that type carry the variant, there should be no association between the variant and the estimated cell type proportion. The power of this test depends on several factors: the proportion of the cell type that carries the mutation, the heteroplasmy level (i.e., the fraction of mitochondrial transcripts with the variant), and the background noise level. The full derivation of the statistical model is provided in Methods.

When the co-enrichment test yields a significant p-value, we confidently assign the variant to the associated cell type. However, an insignificant result can arise either because the variant is truly absent from that cell type (i.e., the null is true) or due to insufficient statistical power. To assess confidence in a negative result (i.e., that a cell type does not carry a given variant), SUMMIT includes a power analysis in module 2. For each non-significant cell type, SUMMIT simulates the probability of detecting the variant across a range of effect sizes.The effect size represents the strength of association between the variant and cell type, □ncurporating both the Proportion of cells carrying the variant and the heteroplasmy level (fraction of mitochondrial transcripts with the variant) compared to background signal (details in Methods). If the power analysis shows high probability of detection (>80%s) at a given effect size, but the actual test remains non-significant, we can confidently conclude the variant is absent from that cell type at that effect size level. Conversely, if power is low, the non-significant result may simply reflect insufficient data rather than true absence.

As we will show in subsequent examples, rare clones can be detected in spatial data if they are spatially localized together. To improve sensitivity in such cases, we conduct a spot-level co-enrichment test when the global co-enrichment test is inconclusive (Fig. 2B, module 3). This test applies a spatial smoothing kernel to borrow strength from neighboring spots and performs multiple testing corrections across the spatial field. This localized test increases statistical power for detecting variants that are spatially restricted to small, coherent regions (Methods).

For the significant variants identified from SUMMIT’s computational pipelines, we want to describe whether the variants are spatially intermixed or separated from each other, and whether each variant is spatially randomly scattered, or spatially clustered (details in Methods, results in Table 2). For a pair of variants, we designed the spatial intermixing index based on Local Inverse Simpson’s Index (LISI) score ^25^. LISI close to 1 indicates that a pair of variants is spatially separated, while LISI close to 2 indicates that the pair is spatially intermixed. For each variant, we also designed the spatial clustering index based on Moran’s I ^26^, which measures the spatial autocorrelation of the variant allele frequency. A significant p-value from the Moran’s test would suggest that a variant is spatially clustered.

### Validation of computational pipeline in the mouse cell line mixture experiment

As an initial proof-of-concept, we applied the computational pipeline to the mouse xenograft experiment. We observed that out of the 6 mitochondrial mutations co-detected by SUMMIT and MAESTER (Fig. S2C-D), variant 3010_G>A is significantly associated with SW1990 (p=2.39e-67), but not to Capan-2. Variants 9545_A>G, 11152_T>C, 10176_G>A, and 13500_T>C are significantly associated with Capan-2 (p=1.84e-29, 4.36e-10, 2.45e-10, 1.41e-4, respectively). Variant 8002_C>T is detected by MAESTER and mtDNA sequencing, but is not significant in SUMMIT data by the global co-enrichment test. The validation of the spatially mixed and cell line specific mutations 3010_G>A and 9545_A>G demonstrates the utility of the global co-enrichment test (Fig. 2C, top). The power analysis indicates that for all mutations (with the exception of 11152_T>C), the non-significant associations with the other cell line and with murine cell types are not explainable by lack of coverage, giving confidence that these variants are indeed not carried by these cell types (Fig. 2C, bottom).

### Identify spatial mitochondrial clones with cell type-aware significance in simulated data

To evaluate how our global and spot-level tests are affected by different sequencing and biological factors, we synthesized two datasets (Fig. 2D, Fig. S2B, Methods). Dataset 1 was synthesized on an artificial 30 × 30 grid, with 3 clones and each clone carrying 5 variants. We considered different levels of variant allele frequency (VAF), clone size, sequencing coverage, and cell type proportion within each spot. For a given coverage and variant allele frequency, the alternative allele count was sampled from a binomial distribution. The cell type ratio was sampled from a beta distribution. This setup allowed us to flexibly explore how different parameters affect detection precision and recall. In this simulation, variant 1 is associated with cell type 1, variant 2 with cell type 2, and variant 3 with cell type 3.

To better reflect real biological structures and sequencing coverage distributions of SUMMIT, dataset 2 was synthesized based on real SUMMIT data. We explored varying levels of variant allele frequency, clone size and the clonal spread in space. Higher spread means that the spots belonging to the clone are more spread out from the center of the clone. In this simulation, all 3 variants are associated with the carrier cell type (red) but not with the background cell types (gray).

For both synthetic data sets, the minimum p-value of each cell type for variant 1 from the spot-wise test are shown in Fig. 2E (axis is -log10 transformed, so higher bars indicate higher significance). We successfully identified variant 1 to be significantly coenriched with expected cell type, and not with the other cell types, which agrees with the ground truth in both datasets (Fig. 2E). As expected, the accuracy of the co-enrichment test, in terms of the F1 score, increases with increased clone size, VAF, and coverage, and decreases with lineage spread in space (Fig. 2F). These in silico experiments offer guidance with regards to the coverage needed for reliable clone detection.

### SUMMIT reveals spatially-segregated clones in Barrett’s esophagus

After validating SUMMIT’s experimental and analytical workflows in the mouse xenograft model system and through simulations, we next assessed its performance on human samples. We selected a nondysplastic Barrett’s esophagus sample as our first test case, as we had previously characterized this tissue type using scRNA-seq and mitochondrial lineage tracing^27^. In Barrett’s esophagus, the normal stratified squamous epithelium is replaced by intestinal-like columnar epithelium containing crypts with goblet cells, foveolar cells, enterocytes, enteroendocrine cells, and intestinal stem cell populations^28,29^. Prior studies have established that Barrett’s esophagus lesions harbor complex clonal and subclonal architecture^18,27,30^, but the spatial distribution of clones within these lesions remains poorly understood.

We performed SUMMIT on a pinch biopsy from a patient with nondysplastic Barrett’s esophagus, collected during routine surveillance endoscopy, to generate spatial transcriptomics and mitochondrial libraries (Fig. 3A-B). From an adjacent tissue slice, we extracted genomic DNA to confirm the presence of variants through mtDNA sequencing (Fig. 3C). For spatial transcriptomics analysis, we first applied RCTD using a single cell sequencing atlas^27^ containing both epithelial (esophageal squamous and Barrett’s esophagus) and non-epithelial (vascular, immune, and stromal) populations (Fig. S3B). Cell type deconvolution revealed a predominantly metaplastic epithelium, with a region of squamous tissue localized to the top of the array (Fig. 3B, Fig. S3C). Gene expression analysis identified differentiated intestinal-like cell types commonly associated with Barrett’s esophagus, including enterocytes, goblet cells, and an ***OLFM4***^***+***^ progenitor population (Fig. S3A) ^31,32^.

**Figure 3:**
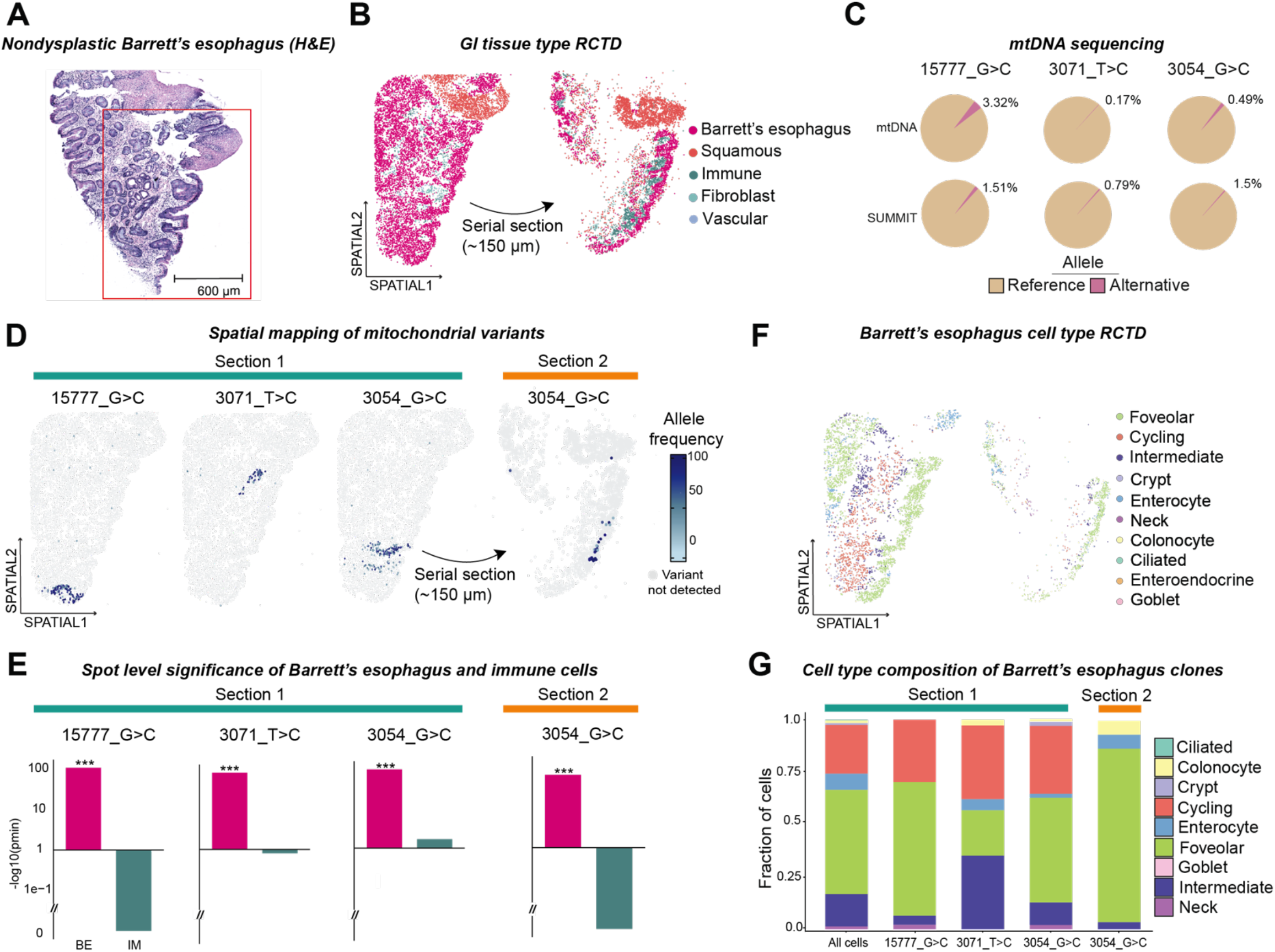
Spatially segregated clones in Barrett’s esophagus. (A) H&E staining of nondysplastic Barrett’s esophagus biopsy showing characteristic glandular architecture with visible crypts. Scale bar: 600 μm. (B) Cell type annotation using RCTD with published Barrett’s esophagus reference atlas^27^. Spatial mapping reveals predominantly Barrett’s esophagus epithelium with a squamous patch as well as immune, fibroblast, and vascular populations in both serial sections. (C) mtDNA sequencing validates the heteroplasmy frequencies for three variants (15777_G>C, 3071_T>C, 3054_G>C) detected by SUMMIT. (D) Spatial mapping of mitochondrial variants reveals three spatially confined clones, each restricted to distinct tissue regions. Section 2 shows presence of 3054_G>C in the same tissue region. (E) Spot-level significance testing confirms significant association of all three variants with Barrett’s esophagus epithelium (BE) but not with immune cells (IM) in both tissue sections. (F) Detailed cell type deconvolution showing spatial organization of specialized Barrett’s esophagus cell types. (G) Cell type composition analysis of each clone showing representation of multiple Barrett’s esophagus cell types within individual clones, consistent with crypt-based organization spanning from base to lumen.

We then performed variant calling on the mitochondrial enriched library, which provided approximately 10-fold greater coverage of mitochondrial transcripts (Fig. S3E). To identify true somatic variants, we filtered out commonly occurring population variants using the MITOMAP database^33,34^, as well as potential nuclear mitochondrial DNA artifacts^35^. This filtering approach yielded heteroplasmic mitochondrial variants in the nondysplastic Barrett’s esophagus sample. We next projected the allele frequencies of each validated variant onto matched spatial coordinates to map the organization of clonal populations within the tissue. Spatial mapping reveals three spatially-confined clones (15777_G>C, 3054_G>C, and 3071_T>C), each confined to a distinct region of the tissue (Fig. 3D). To investigate the depth of these crypt-localized variants, we profiled a distant serial section (∼150 μm) from the same biopsy and found evidence of one spatial variant, 3054_G>C, in the same tissue region (Fig. 3D). The other two spatial variants were not detected in this distant serial section.

We next sought to test whether these mitochondrial variants were significantly enriched in the regions of the tissue containing Barrett’s esophagus cell types. We used our spot-level cell type significance test to evaluate the co-enrichment between each variant (15777_G>C, 3054_G>C, and 3071_T>C) and Barrett’s esophagus or immune populations. For each variant in the original slice and the distant serial section, we found highly significant co-enrichment between the mtDNA variant and Barrett’s esophagus expression program (spot-level minimum p-value=5.15e-109, 1.07e-74, and 3.44e-60, respectively). Conversely, there was no significant co-enrichment between these variants and non-Barrett’s cell types, despite having sufficient coverage for each variant in the non-Barrett’s cell types to ensure adequate power (Fig. S3F). This confirms that these spatially-confined variants are restricted to Barrett’s esophagus cell types (Fig. 3E).

Since the Barrett’s esophagus epithelium contains multiple specialized cell types, we performed a second cell type deconvolution using a detailed Barrett’s esophagus reference dataset to achieve finer cell type resolution (Fig. S3D). From this reference, we found foveolar cells localizing at the tissue lumen, intermediate cells below the lumen, and cycling cells distributed towards the base of the gland (Fig. 3F). Examining the spatial distribution of our three variants revealed a pattern that each clone appeared to extend vertically through the tissue, spanning from basal regions toward the lumen (Fig. 3D). Particularly for variants 15777_G>C and 3054_G>C, we noted that the cell type composition of each captured the full base-to-lumen composition observed in these Barrett’s glands (Fig. 3G). In 3071_T>C, we appear to have captured a cross-section of a crypt, containing enrichment for more of the intermediate cell states. In Section 2 from this block, we sampled regions containing mostly foveolar cells, as these were over represented in the 3054_G>C variant within this section. The diversity of Barrett’s esophagus cell types per clone is consistent with our prior single cell-based work finding multiple mitochondrial clones are present in Barrett’s biopsies containing the full array of cell types^27^. Further, using the spatial information and adjacent H&E, the positions of these clones appear to be restricted to individual crypts. This compartmentalization suggests that a common ancestor within the crypt contained the variant and gave rise to all the progeny within that specific crypt of the tissue.

### Independent clones captured in the normal small bowel epithelium

To test SUMMIT on a different tissue architecture, we collected a normal small bowel sample from a patient with Lynch syndrome during bowel resection. H&E staining of this sample showed intact intestinal architecture with visible villi and crypts (Fig. 4A). In parallel, we generated a paired single nucleus sequencing (snRNA-seq) reference from an adjacent cryopreserved sample for cell type annotation with RCTD (Fig. S4A).

**Figure 4:**
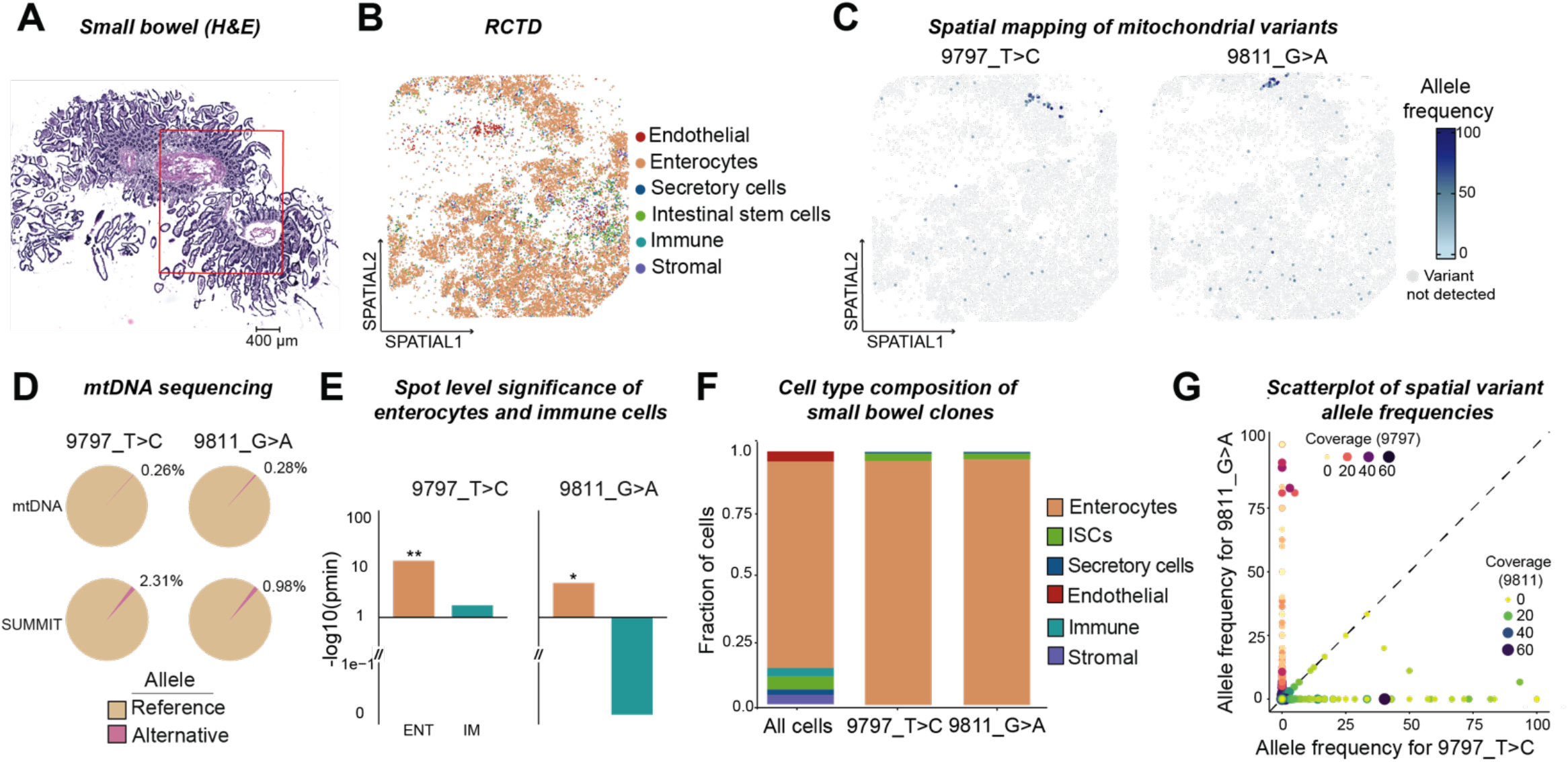
Independent clones captured in the normal small bowel epithelium. (A) H&E staining of small bowel tissue showing intact intestinal architecture with visible villi and crypts. Scale bar: 400 μm. (B) RCTD cell type annotation using paired single-nucleus RNA sequencing reference shows spatial distribution of enterocytes, secretory cells, intestinal stem cells, immune, endothelial, and stromal populations. (C) Spatial mapping of mitochondrial variants 9797_T>C and 9811_G>A, showing adjacent but non-overlapping localization within the intestinal epithelium. (D) mtDNA sequencing detects both variants with heteroplasmy frequencies of 2.36s and 2.39s, respectively (E) Spot-level significance testing demonstrates significant association of both variants with enterocytes (ENT) but not with immune cells (IM). (F) Cell type composition analysis showing both clones are predominantly composed of enterocytes with small populations of intestinal stem cells and secretory cells. (G) Scatterplot of variant allele frequencies for 9797_T>C and 9811_G>A demonstrating independence of the two variants, with high-confidence spots for one variant showing low signal for the other.

Deconvolution of the spatial data revealed that enterocytes comprised the majority of the tissue (81.7%), with smaller populations of endothelial, secretory, stromal, and intestinal stem cells (Fig. 4B, Fig. S4C). In the mitochondrial genotyping data, we detected two spatially confined clones, 9797_T>C and 9811_G>A, both of which were also confirmed in mtDNA from an adjacent slice (Fig. 4C-D, Fig. S4B). While neither variant was significant in any cell type using the global test (Fig. S4E), this was expected given that both variants were restricted to a small region of the tissue section. Using our spot-level cell type significance test, both variants were significantly associated with the intestinal epithelial cells (spot-level minimum p-value=2.99e-15 and 7.94e-06, respectively). Specifically, both variants were significantly associated with enterocytes but not with immune cells (Fig. 4E,F, Fig. S4D). Furthermore, power analysis confirmed that we had sufficient coverage to detect these variants in immune cells if they were present (Fig. S4E).

The alternative allele frequencies of the two variants are very similar in bulk whole-exome sequencing (WES), such that bulk-level subclone identification tools would likely group them into the same cluster^36^. However, spatial projection of allele frequencies reveal variants 9797_T>C and 9811_G>A localize to nearby but non-identical regions of the tissue. To quantify their spatial segregation, we used the Local Inverse Simpson’s Index (LISI), which measures the average number of distinct mutations among each cell’s *k*-nearest neighboring cells in the physical 2-D space (details of LISI score in Methods). The LISI score for variants 9797_T>C and 9811_G>A was 1.08, indicating complete spatial segregation of the two variants. This is further supported by a pairwise scatterplot of the allele frequencies of the two variants, with the points sized and colored by the coverage of the variant with lesser VAF (Fig. 4G). This visualization revealed that spots with high confidence for one variant (both high coverage and allele frequency) are mostly confident non-carriers for the other variant (high coverage but low allele frequency). Spots appearing to contain both variants had either low coverage or low allele frequency for at least one variant. This confirms that variants 9797_T>C and 9811_G>A represent two independent clones that co-occur in adjacent regions of the tissue.

### Subclonal relationships can be inferred within normal gastric epithelium

We next tested SUMMIT on normal gastric cardia collected during routine endoscopy. H&E staining showed intact gastric glands with the characteristic pit-to-base architecture (Fig. 5A). We used scRNA-seq data from Gier et al.^27^ for cell type deconvolution (Fig. S5A). Cell type annotation of the spatial data revealed the expected spatial organization of gastric epithelium: pit cells positioned at the lumen, chief cells at the gland base, cycling and parietal cells distributed throughout the middle regions, and scattered cycling and enteroendocrine cells present throughout the tissue (Fig. 5B, Fig. S5D).

**Figure 5:**
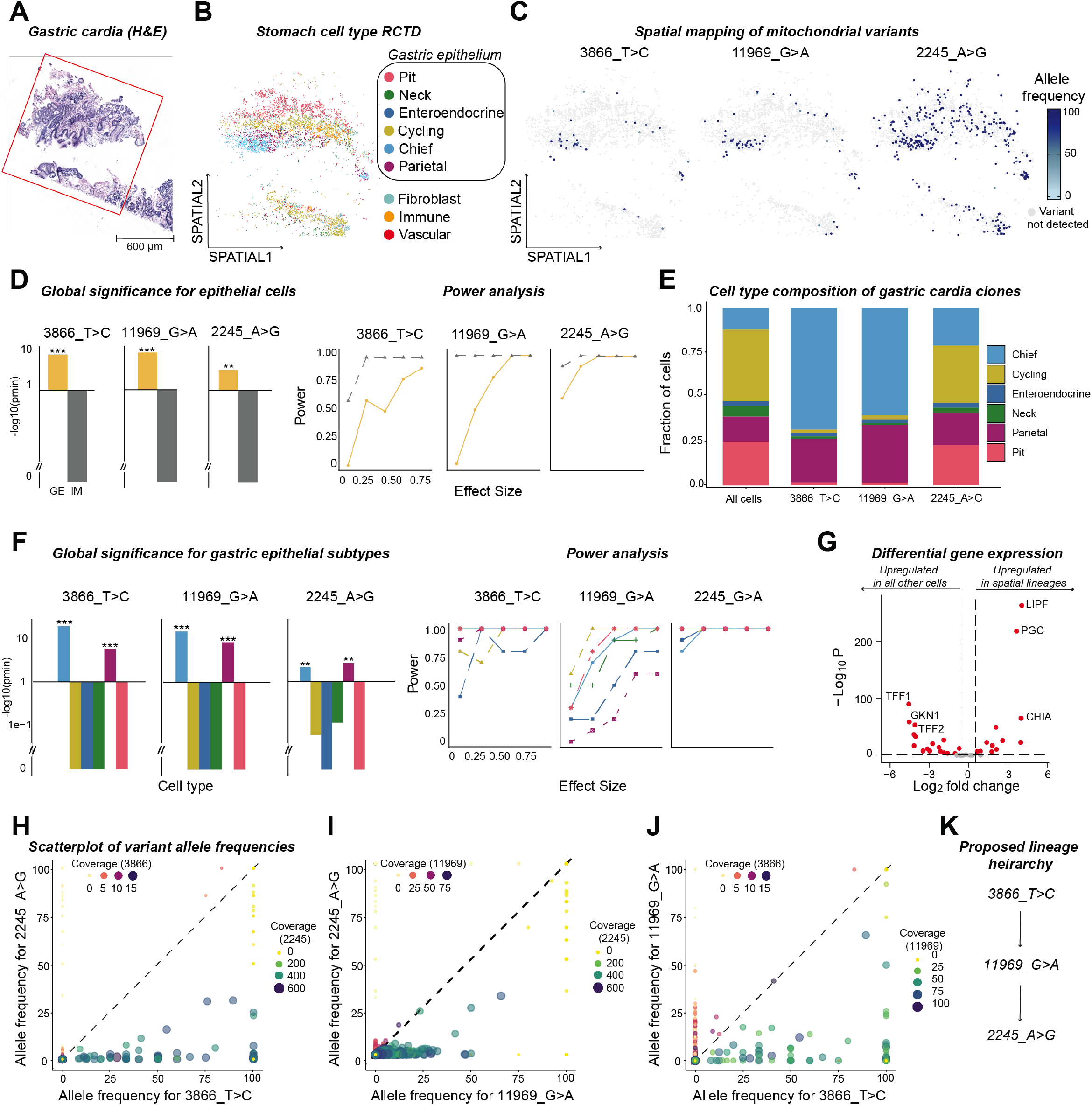
Subclonal relationships can be inferred within normal gastric epithelium. (A) H&E staining of normal gastric cardia showing intact gastric glands with characteristic pit-to-base architecture. Scale bar: 600 μm. (B) Cell type annotation with RCTD showing localization of gastric epithelial subtypes (pit, neck, enteroendocrine, cycling, chief, and parietal cells), along with stromal and immune populations. (C) Spatial mapping of mitochondrial variants 3866_T>C, 11969_G>A, and 2245_A>G, showing varying degrees of intermixing. (D) Global significance testing demonstrates significant association of all three variants with gastric epithelium (GE) but not with immune cells (IM). Power analysis confirms adequate coverage for confident cell type assignment. (E) Cell type composition analysis showing that spatially-restricted clones (3866_T>C and 11969_G>A) are predominantly composed of chief cells and parietal cells, reflecting their basal gland location. Global significance testing confirming significant enrichment of variants 3866_T>C and 11969_G>A in chief and parietal cells across gastric epithelial subtypes. (G) Differential gene expression analysis showing enrichment of chief cell markers *LIPF* and *PGC* in spatially-restricted clones. (H-J) Pairwise scatterplot analysis of variant allele frequencies revealing subclonal relationships;2245_A>G appears as a subclone of both 3866_T>C and 11969_G>A, while 11969_G>A appears as a subclone of 3866_T>C. (K) Proposed lineage hierarchy showing inferred evolutionary relationships between the three variants based on co-occurrence patterns.

In this sample, we identified three mitochondrial variants, including two spatially distinct variants, 3866_T>C and 11969_G>A, and one diffuse variant, 2245_A>G (Fig. 5C, Fig. S5B-C). All three variants are spatially intermixed, with pairwise LISI scores equal to 2 for variant pair 2245_A>G and 11969_G>A, 1.88 for 2245_A>G and 3866_T>C, and 1.90 for 11969_G>A and 3866_T>C (Table 1).

Using our global cell type co-enrichment test, all three clones were significantly associated with epithelial cell types (p=1.84e-6 for 3866_T>C, 1.55e-09 11969_G>A, 1.2e-03 for 2245_A>G), and showed no co-enrichment with non-epithelial cell types despite having sufficient coverage and power (Fig. 5D). Looking at intra-clonal cell type diversity, cell type composition of the spatially restricted clones reflected their anatomical location. Clones 3866_T>C and 11969_G>A consisted primarily of chief cells (68.75% and 60.55%, respectively) and parietal cells (24.7% and 32.6%, respectively). These findings are consistent with the spatial positioning of these variants being restricted to the gland base region where these cell types predominate (Fig. 5E). Testing global associations with specific epithelial subtypes confirmed that variants 3866_T>C, 11969_G>A, and 2245_G>A were significantly enriched in both chief cells (p=2.02e-20, 1.2e-15, 7.2e-03) and parietal cells (p=1.44e-11, 1.20e-15, 2.2e-03) (Fig. 5F, Fig. S5E-F).

We next performed differential gene expression analysis to identify transcriptional differences between epithelial cells contained within clones 3866_T>C and 11969_G>A and all other epithelial cells. Differential gene expression revealed enrichment for chief cell markers, *LIPF* (log2FC = 4.05, p < 1e-6) and *PGC* (log2FC = 3.65 p < 1e-6), in the spatially-restricted clones (Fig. 5G). The enrichment of chief cell markers corresponded with the spatial localization of these clones at gland bases where *LIPF* expression was highest, confirming the chief cell identity of these clones (Fig. S5G).

Since these three variants exhibit spatial intermingling, we used pairwise scatterplots to visualize their co-segregation. First, consider the relationship between 3866_T>C and 2245_A>G (Fig. 5H).There are 5 spots that confidently carry both variants (coverage > 5, VAF > 25), while there are 35 spots that confidently carry 3866_T>C (coverage > 5, VAF > 25) but not 2245_A>G (coverage > 5, VAF < 25).Conversely, there are also 56 spots that carry 2245_A>G but have low VAF for 3866_T>C. However, these cells that carry 2245_A>G but not 3866_T>C have insufficient coverage of 3866_T>C, and thus the low or zero VAF could be due to drop-out. Based on this, we infer that cells carrying 2245_A>G form a subclone of cells that carry 3866_T>C.

Now consider the relationship between 11969_G>A and 2245_A>G (Fig. 5I). As above, some spots carry both 2245_A>G and 11969_G>A in high confidence (coverage> 5, VAF>25), while others carry 11969_G>A but not 2245_A>G. Conversely, all spots carrying 2245_A>G and no detectable 11969_G>A have extremely low coverage for 11969_G>A, and thus the low or zero VAF for 11969_G>A in these spots could be due to drop-out. This implies that 2245_A>G is likely subclonal to 11969_G>A.

Finally, consider the relationship between 11969_G>A and 3866_T>C (Fig. 5J). There are 5 spots that are high confidence carriers of both variants, and there are 53 spots that are high confidence carriers of 3866_T>C but not 11969_G>A. In contrast, there is only one spot carrying 11969_G>A without also carrying 3866_T>C, as the spots with 11969_G>A but low or zero VAF for 3866_T>C had insufficient coverage. This suggests that 11969_G>A is a subclonal mutation within the 3866_T>C clone.

Taken together, these relationships imply that cells carrying either 11969_G>A or 3866_T>C should also carry 2245_A>G, but not vice versa (Fig. 5K). This inference is counterintuitive given the spatial distribution of 2245_A>G, which appears more diffuse than either 3866_T>C or 11969_G>A. The paradox is resolved when visualizing the coverage maps: coverage for 3866_T>C and 11969_G>A is uneven and concentrated in the gland base, whereas coverage for 2245_A>G is broader and more uniform (Fig. S5G). This example demonstrates that SUMMIT can reveal subclonal relationships within tissues. However, accurate inference requires consideration of variant-specific coverage.

### Malignant clones exhibit spatial intermixing in colorectal cancer

In addition to normal and premalignant tissues, we also applied SUMMIT to a colorectal cancer sample to assess its performance in neoplastic tissue with higher mutational burden (Fig. 6A). The tissue consisted predominantly of malignant epithelium with areas of infiltrating stroma (Fig. 6B, Fig. S6A-B,D). SUMMIT identified six mitochondrial variants, five of which significantly associated with the malignant epithelial compartment and not with non-epithelial cell types, confirming their specificity as cancer cell markers (Fig. 6C-D, Fig. S6E, Table 1). All six variants were also Independently validated by bulk mtDNA sequencing (Fig. 6E).

**Figure 6:**
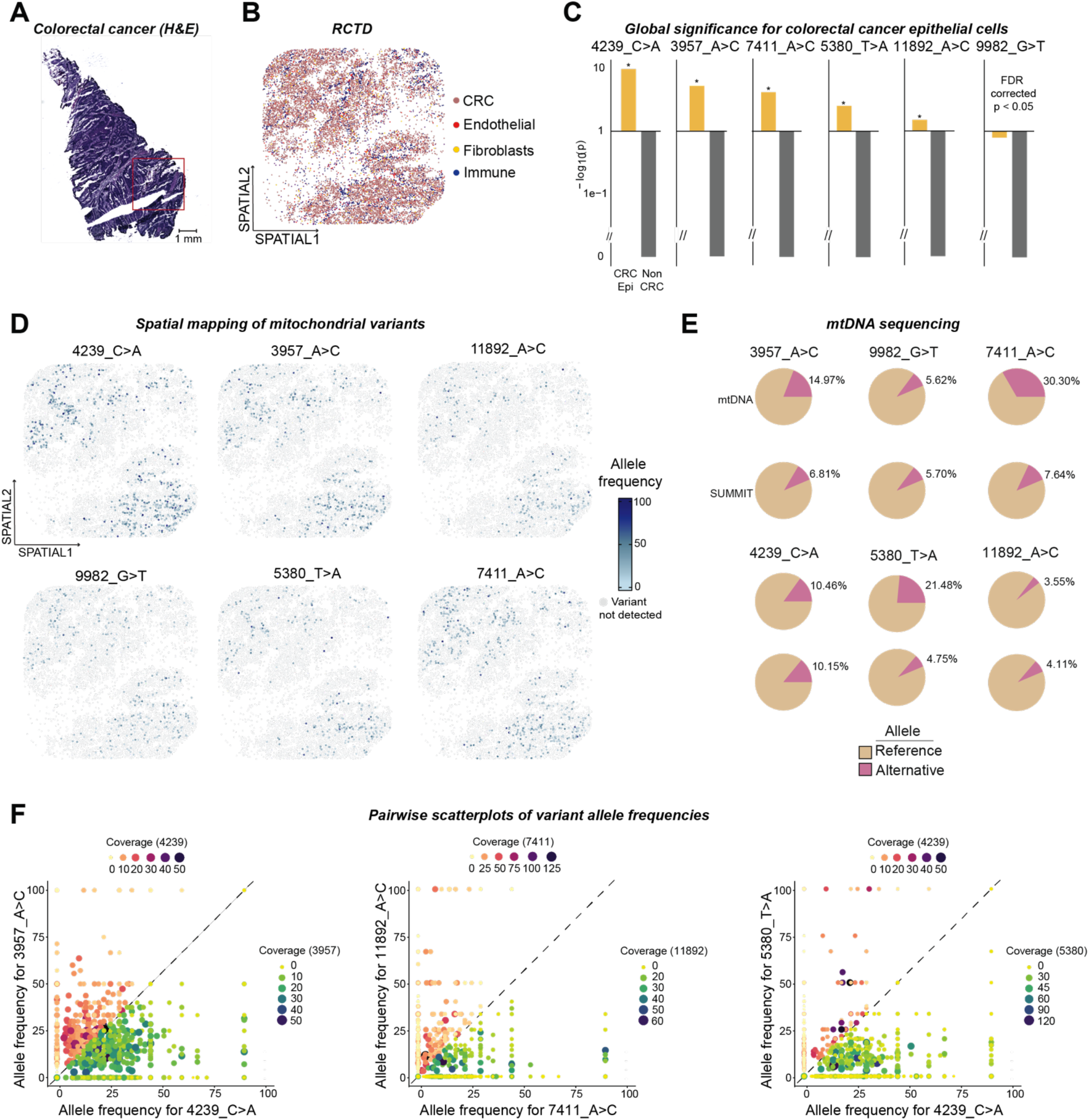
Malignant clones exhibit spatial intermixing in colorectal cancer. (A) HE staining of colorectal cancer sample showing malignant epithelial architecture with disrupted tissue organization. Scale bar: 1 mm. (B) Cell type annotation with RCTD revealing predominantly colorectal cancer (CRC) epithelium with infiltrating endothelial, fibroblast, and immune populations. (C) Global significance testing for six mitochondrial variants (4239_C>A, 3957_A>C, 7411_A>C, 5380_T>A, 11892_A>C, 9982_G>T) showing significant association with colorectal cancer epithelium (CRC Epi) but not with non-epithelial cell types (Non CRC). Variant 9982_G>T is significant with FDR correction for multiple testing. (D) Spatial mapping of all six mitochondrial variants showing spatial intermixing across the malignant tissue. (E) mtDNA sequencing validating the presence of all six variants. (F) Pairwise scatterplot analysis of variant allele frequencies showing evidence of co-occurrence for variant pairs 3957_A>C – 4239_C>A, 11892_A>C – 7411_A>C, and 5380_T>A– 4239_C>A. Spots carrying high-confidence signal for one variant generally lack coverage for others, suggesting potential common clonal origin of all detected variants.

Unlike the spatially restricted clones observed in normal and premalignant tissues, these cancer-associated variants exhibited broad spatial overlap and intermixing across the tissue. This spatial pattern is consistent with the clonal expansion and architectural disruption characteristic of malignant progression. Quantification using the LISI showed high mixing of variants within local neighborhoods (LISI score of all pairs close to 2) indicating the absence of spatial segregation (Table 2).

To determine whether these variants marked independent clones or a single clonal population, we analyzed their pairwise co-occurrence. For every pair of variants, we found spots that carried both with high confidence. In cases where a spot appeared to carry only one variant, the other variant always had insufficient coverage (Fig. 6F). This pattern indicates that we can not rule out the configuration where all six mutations co-occur within a single clonal population.

Together, these findings support a model in which the malignant epithelium is derived from a common progenitor harboring multiple mitochondrial mutations, followed by clonal outgrowth and expansion. This is in stark contrast to the spatially restricted, crypt-confined clones observed in nondysplastic Barrett’s esophagus, normal gastric cardia, and small bowel tissue, further underscoring how spatial clonal architecture shifts in the transition to neoplasia.

## Discussion

Here, we developed and validated SUMMIT as a method for resolving clonal lineages in intact tissues using endogenous mitochondrial mutations. By integrating mitochondrial lineage tracing with spatial transcriptomics, SUMMIT enables joint interrogation of how cellular ancestry and spatial organization contribute to tissue architecture across diverse tissue types.

Our analysis of normal and premalignant tissues revealed fundamental principles of tissue organization. Across the samples we analyzed, we observed that clones maintain spatial coherence, with individual clones confined to discrete anatomical units. In Barrett’s esophagus, clones respected crypt boundaries and contained the full complement of differentiated cell types expected along the crypt-lumen axis. This base-to-lumen organization (encompassing stem, progenitor, and differentiated cells within single clones) provides direct evidence that Barrett’s crypts contain contiguous clones that span the full differentiation hierarchy — a finding that parallels established intestinal biology^37,38^.

In the small bowel, we identified adjacent but non-overlapping clones occupying the same tissue region, suggesting derivation from neighboring stem cell niches. In contrast, the gastric sample showed evidence of ongoing clonal evolution, with a nested configuration between the 3 identified variants demonstrating that SUMMIT can capture evolutionary relationships between clones. Finally, the colorectal cancer sample revealed a different clonal architecture, with six mitochondrial variants showing complete spatial intermixing rather than the compartmentalized organization observed in normal tissues. This pattern may reflect the advanced stage of malignancy and clonal expansion that occurred prior to sampling, though the relatively small tissue area analyzed limits our ability to assess broader tumor heterogeneity that might exist at larger spatial scales.

A key innovation of SUMMIT is the statistical framework for assigning variants to specific cell types within spatially mixed spots. This is critical because, even at 10μm resolution, spots frequently contain multiple cell types. SUMMIT combines cell type deconvolution with lineage-specific co-enrichment testing and power analysis, ensuring that lack of significant co-enrichment reflects true absence rather than insufficient power. To improve power for small clones that are spatially concentrated, SUMMIT employs an efficient spot-level scan statistic rather than a global tissue-wide test. This flexible but rigorous framework allowed us to confidently determine the cell type of origin of the detected clones, even for rare clones that are restricted to specific tissue niches. This was demonstrated on a variety of tissue types harboring diverse clonal configurations, both diffuse and spatially restricted.

SUMMIT advances beyond existing mitochondrial lineage tracing approaches in several key ways. The added spatial context builds upon existing single cell RNA and multiomic platforms that are receiving wide adoption. While imaging-based methods like cytochrome c oxidase staining provide spatial resolution^19,39^, they are limited to specific mitochondrial mutations affecting enzyme function. Copy number-based lineage tracing yields limited clonal resolution on tissues that maintain genomic stability^40,41^. In contrast, SUMMIT leverages the entire mitochondrial genome as a source of lineage markers while simultaneously capturing full transcriptomic information.

The current implementation of SUMMIT has important limitations. It requires fresh-frozen tissue, limiting its applicability to the vast archives of FFPE clinical samples. Future efforts may build on recent innovations, such as Patho-DBiT, which enables capture from fragmented RNA via polyadenylation strategies^42^. Another limitation is capture efficiency. Although mitochondrial transcripts are significantly enriched in the SUMMIT libraries, coverage remains uneven across the mitochondrial genome. Future strategies could draw from approaches developed for scRNA-seq that enable more efficient direct capture from mitochondrial DNA^43^. Broadly, as spatial technologies improve capture efficiency, SUMMIT’s performance will improve in parallel. In addition, the principles underlying SUMMIT can be extended to other spatial profiling platforms, including those with higher resolution or multiomic capability^44–46^, broadening the scope and utility of this approach.

Overall, this work presents SUMMIT as the first technology providing unbiased spatial mitochondrial clone tracing with spatial transcriptomics. This unique capability enables researchers to dissect how cellular ancestry and spatial context jointly drive tissue heterogeneity. The technology’s applications span normal tissue development and disease progression, offering insights on the clonal architecture at all stages of clinical disease development in patient tissues.

## Methods

### Primary sample collection and cryobanking

Primary gastrointestinal tissue was obtained following written informed consent under IRB approval at the Hospital of the University of Pennsylvania (Protocol #813841). Barrett’s esophagus and gastric cardia tissues were collected via pinch biopsies during routine endoscopy from a previously diagnosed patient (G.W.F). Small bowel tissue was obtained from bowel resection of a patient previously diagnosed with Lynch syndrome (E.E.F). Colorectal cancer sample was obtained during surgical removal of cancerous region (B.W.K). Samples were immediately washed in ice-cold Dulbecco’s phosphate buffered saline (DPBS; Corning, 21-031-CV), embedded in optimal cutting temperature solution (OCT) (Fisher, 23730571) and snap-frozen in an isopentane-dry ice bath (O35514, Fisher) and stored at -80°C.

### Subcutaneous injection for tumor generation

A 2:1 mixture of human pancreatic adenocarcinoma cell lines, Capan-2 (HTB-80™, ATCC®, Manassas, VA, USA) and SW1990 (CRL-2172™, ATCC®, Manassas, VA, USA) was subcutaneously injected into both flanks of an immunocompromised, NSG mouse (total 1.5 × 10^6^ cells). Tumors grew for 17 days until the volumes averaged 90 to 100 mm^3^ by caliper measurement. The mouse was sacrificed and tumors were harvested and placed in ice-cold DPBS. One tumor was embedded in OCT for spatial transcriptomic analysis and the other immediately was dissociated for single-cell RNA sequencing.

### Isolation of single cells from murine tumor

Murine tumor was minced using dissection scissors and resuspended in 500 μL DPBS containing Liberase TH (1 WU/mL, Roche, 5401151001) and RQ1 DNase (0.5 U/mL, Promega, M6101). Tissue was digested in a thermomixer (37°C, 400 RPM) for 10 minutes with trituration every 5 minutes. After 10 minutes, the solution was briefly centrifuged and supernatant was filtered through a 70 µm filter and collected in 8mL DPBS + 1s BSA (Miltenyi Biotec, 130-091-376) on ice. Large tissue pieces were resuspended in 500 μL fresh enzyme-PBS solution and digested a total of three times. Single-cell suspension was filtered through a 35 µm filter and centrifuged for 4 minutes (400 x g, 4°C). Supernatant was removed and the pellet was washed with 1mL of cold DPBS + 0.04% BSA twice. Cells were counted and assessed for viability using Trypan blue (Gibco, 15250061). Approximately 15,000 cells were loaded onto the 10x Chromium Controller for GEM generation (kit: 1000688, beads: 10000692, chip: 10000690).

### Isolation of single nuclei from patient tissue

Cryoembedded tissue was thawed in DPBS to remove excess OCT, washed with cold DPBS, and resuspended in 1 mL TST buffer. TST buffer contains 146 nM NaCl (Sigma Aldrich, 59222C), 10 mM Tris-HCl (pH 7.4, NEB BioLabs, E7496AA), 1mM CaCl_2_ (Sigma Aldrich, C5670-100g), 21 nM MgCl_2_ (Invitrogen, AM9530G), 0.03% Tween 20 (Life technologies, 28320), 0.01% BSA, 1:100 RNAase inhibitor (Sigma Aldrich, 3335402001) in H2O. Tissue was minced for 10 minutes on ice. Minced nuclei were filtered through a 50 µm filter into a 50 mL conical and washed with 1mL of TST buffer. Solution was diluted with 3 mL ST buffer (146 nM NaCl, 20 nM Tricine (Thermo Fisher, 42020132-1), 1mM CaCl_2_, and 21 nM MgCl_2_ in H2O) and transferred into a 15 mL conical. Nuclei solution was centrifuged at 500g for 5 minutes (brake setting 5) at RT. Pellet was resuspended in 200 μL nuclei resuspension buffer (1% BSA and 1:40 RNAse inhibitor in PBS). Diluted solution was filtered through a 35 µm filter and counted in trypan blue with a hemocytometer. Approximately 15,000 nuclei were loaded onto the 10x Chromium Controller for GEM generation.

### 10x Genomics Chromium GEM-X Single Cell 3’ v4 library preparation and paired-end sequencing

Single-cell and single-nucleus suspensions were loaded onto a 10x Chromium Controller for GEM generation. Dual index libraries were generated following the GEM-X Single Cell 3′ v4 library preparation protocol (part no. 1000689) and quantified using Agilent 2100 Bioanalyzer with High Sensitivity DNA Kit (Agilent, 5067-4626). Indexed libraries were sequenced on either the Illumina NextSeq 550 (10 cycles per index, 28 cycles Read 1, 43 cycles Read 2) or Element AVITI (10 cycles per index, 28 cycles Read 1, 110 cycles Read 2). Single cell RNA-sequencing of the murine tumor averaged >18,000 reads per cell. Single nucleus RNA sequencing of colorectal cancer and small bowel averaged >10,000 reads per nucleus.

### Genome alignment and count matrix generation with Cell Ranger v8.0

Raw base call files from Illumina or Element were demultiplexed to generate FASTQ files using bcl2fastq (v2.20.0.422) or bases2fastq, respectively. Fastq files were aligned to the GRCh38-2024-A genome or combined human/mouse genome GRCh38-GRCm39-2024-A using CellRanger (v8.0) to generate count matrices.

### scRNA-seq dimensionality reduction, clustering, and cell type annotation in Seurat

Count matrix (.h5) was converted to a Seurat object (v5.1.0) and filtered to retain cells with >300 genes, >250 transcript counts, and <10% mitochondrial reads. Doublets were removed using scDblFinder^47^ (v1.18.0) with nfeatures = 3000 and includePCs = 1:20. Following log-normalization and scaling, PCA (2000 variable features), then UMAP (10 components), dimensionality reduction was performed. Low dimensional projection was plotted using FindNeighbors (10 components) and FindClusters (resolution=0.1). Differential genes identified by Wilcoxon rank-sum test were used to annotate Capan-2, SW1990, and mouse populations. The filtered count matrix and cluster annotations were used to generate an RCTD reference using spacexr^21^ (v2.2.1).

### Curio Seeker library preparation and paired end sequencing

Fresh frozen tissue was equilibrated to -20°C for 20 minutes before sectioning. OCT-embedded tissue was mounted and sliced at 10µm thickness with a 10° cutting angle in a cryostat. The section was melted onto a Curio Seeker 3×3 tile by placing a finger on the bottom side of the slide. Tile was removed from the slide and placed in a 1.5mL Eppendorf tube containing Hybridization mix. Samples were prepared according to the 3×3 Curio Seeker protocol. cDNA was amplified from bead-bound transcripts for 10 cycles and 600 pg cDNA was tagmented using the Nextera XT Library Prep Kit (Illumina, FC-131-1024). Dual indexed libraries were sequenced on either the Illumina NextSeq 550 or Element AVITI (8 cycles per index, 50 cycles Read 1, 90 cycles Read 2).

### Genome alignment with Curio Seeker

Raw base call files from Illumina or Element were demultiplexed to generate FASTQ files using bcl2fastq or bases2fastq, respectively. Fastq files were aligned to the GRCh38genome or combined human/mouse genome GRCh38-mm10 using CurioSeeker (v3.0) to generate count matrices.

### Spatial RNA-sequencing dimensionality reduction, clustering, and cell type annotation

Spatial count matrix (.h5ad) was converted to a Seurat object (v5.1.0). Cell type proportions were determined by robust cell type deconvolution (RCTD) using ‘doublet’ mode (min_UMI = 100). Annotations were assigned the cell type that occupied > 75% of the bead. Beads lacking a dominant cell type were removed. Spatial RNA-sequencing of the mouse tumor was filtered to retain beads with log10(UMIs) > 1.5 and nFeature_RNA > 35. Low dimensional projection was plotted using spatial transcriptomic clustering algorithm, Banksy^48^. To run Banksy, 8000 variable features were detected with the “vst” selection method. PCA was run using SCT assay, and low dimensional projection was plotted using FindNeighbors (25 components), FindClusters (resolution=0.25), and RunUMAP(dimensions = 25, n.neighbors = 100, min.dist = .1, spread = 3). Barrett’s esophagus, gastric cardia, small bowel, and colorectal cancer spatial RNA-sequencing data were processed exclusively with RCTD (min_UMI = 35) for cell type annotation. Using Seurat, spatial gene expression plots were generated after log-normalization and scaling using the SpatialFeaturePlot command.

### MAESTER enrichment and sequencing

Mitochondrial transcripts were enriched from the spatial cDNA libraries using the MAESTER^20^ protocol. Full length cDNA underwent 6 additional cycles of amplification before allocation of 10 ng to 12 separate PCR reactions containing primers that tile the mitochondrial transcriptome. cDNA was amplified for 6 additional cycles according to the Curio Seeker protocol, pooled, and purified using Ampure XP (Beckman Coulter, A63881). Sequencing primer binding sites and dual indexes were added through 6 additional PCR cycles. The concentration of final libraries was between 1-5 ng/μL with 800-1,200 bp fragment sizes. Concentrations were determined by Qubit, and library sizes were determined by BioAnalyzer trace. Libraries were sequenced on an Illumina NextSeq 550, Illumina NextSeq 1000/2000, or Element AVITI with a 300-cycle kit (43 cycles for Read 1, 250 cycles for Read 2, and eight cycles for both indices).

### MAESTER sequencing data pre-processing and mapping

FASTQ files were generated from raw base calls using bcl2fastq or bases2fastq using the i7 index. Reads with spatial barcodes not matching a corresponding whitelist of barcodes present in the Curio Seeker tile were removed. R2 reads were processed to include the spatial barcode and UMIs as SAM tags. Reads were aligned using STAR^49^ and filtered to retain only mitochondrial genome alignments using custom scripts.

### Mitochondrial genome variant calling

BAM files with spatial barcodes and UMIs as SAM tags were processed with maegatk to call mitochondrial mutations. maegatk (https://github.com/caleblareau/maegatk) was run with the following arguments:-g rCRS - mb 1 -mr 1.

### Filtering of mitochondrial variants

Allele frequencies, mean coverage, mean Q score, and allele frequency quantiles of mitochondrial variants were calculated from the maegatk object in R. High quality variants were selected using the following parameters: mean coverage of >3, mean quality of >21, and AF of >25% in at least 1s of beads. Variants identified as germline by bulk mtDNA sequencing and mutations present in nuclear mitochondrial DNA were removed from analysis^35^.

High quality variants detected in the single cell RNA-seq dataset of the xenograft tumor were selected using mean coverage of >3, mean quality of >27, and AF of >25% in at least 1% of beads was used to filter. Variants that passed aforementioned filters were plotted onto matched spatial coordinates. Spatial allele frequency plots, generated with custom ggplot code, included beads with a minimum coverage of 3 for the variant of interest and allele frequency >35%. Bulk heteroplasmy estimations were calculated using beads that had an allele frequency >35% for a particular variant.

### Cell type Decompositions of Spatial Transcriptomics Data

For all the spatial transcriptomics data, each spot was decomposed with cell types from a reference dataset by *Robust Cell Type Decomposition (RCTD)* with R package *spacexr v2*.*2*.*1*. For the cell line mixture experiment, paired single cell RNA-sequencing data was used as reference. For the colorectal carcinoma and small bowel disease samples, a paired snRNA data was used as reference. For the Barrett’s Esophagus and gastric cardia samples, the published scRNA-seq atlas in Gier et al.^27^ was used as reference.

Overall, when paired scRNA-seq or scRNA-seq data is not available, publicly available cell atlases can be used as reference datasets for decomposition.

### Global Cell type Test

We describe here the model underlying the statistical test that cell type *k*is coenriched with variant *j*. Co-enrichment means that we expect a proportion of cells from cell type *k*to be carriers for the variant *j*.

For variant *j*, let *θ*_0j_ represent its background VAF, that is, the probability of a read carrying the alternative allele of the variant within a spot that does not carry it. Background VAF quantifies the false positive rate of calling a variant. Let VAF represent the expected VAF within carrier cells, that is, the heteroplasy proportion.

For celltype *k*∈{1,2, …*K*}, we let Π_*jk*_ be the proportion of cells within the cell type that carry variant *j*. Our null hypothesis is thus:

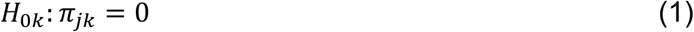

versus the alternative that Π_*jk*_ > 0.

Within spot *s*, let *W*_*sk*_ be the proportion of cell type *k* estimated via deconvolution. We start by defining latent variable *Z*_*sij*_, the allele on read *i* mapping to spot *s* for variant j. *Z*_*sij*_ is unobserved in our data, but has the distribution:

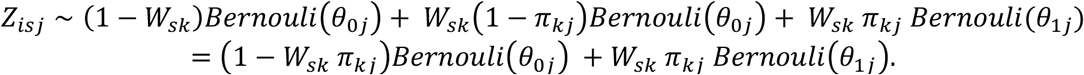

That is, if the read comes from cell type *k*(with probability *W*_*s k*_), then it has probability Π_*j k*_ of being a carrier of the variant. If it were the carrier of the variant, then it has probability *θ*_1*j*_ of containing the alternative allele. If we assume, for the purpose of this hypothesis test, that cell type *k*is the only possible carrier cell type, then in all other cases read *i* would carry the alternative allele with background probability *θ*_0*j*_. Summing up the reads that map to spot *s*, we have the distribution for *X* _*s j*_, alternative allele counts in spot *s*:

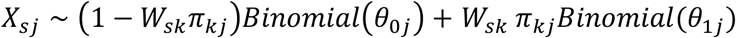

We take the expectation of both sides of the above, and rearrange to get:

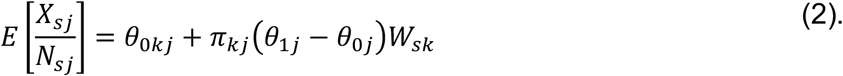

We make the reasonable assumption that *θ*_1*j*_ > *θ*_*0 j*_, that is, the expected carrier VAF is strictly greater than the background VAF, then our null hypothesis (1) is equivalent to the slope Π_*k j*_ (*θ*_1*j*_ *– θ*_*0 j*_) = 0 in the above linear model. We can estimate this slope through a weighted regression of the observed VAFs 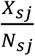 on *W*, weighted by coverage *N*_*sj*_. We say that variant *j* is significantly co-enriched with cell type *k*if the slope in this regression is significantly positive, i.e. p-value less than a preset threshold and positive estimated value.

We performed this regression test for each detected mitochondrial variant, after quality filtering, and controlled the FDR at 0.05 through Benjamini Hochberg procedure.

### Power Analysis

For a given cell type *k*, if the global cell type test does not yield a significant test result it may be due either to the fact that cell type *k* truly does not carry the variant, or to low sequencing coverage and signal dropout. To figure out whether a cell type is truly not a carrier, we designed a corresponding power analysis.

Our model assumes that the observed alternative allele counts *X*_*sj*_ of variant *j* and spot *s* follows a binomial distribution with coverage *N*_*sj*_ and the probability of sampling the alternative allele reads equal to the variant allele frequency *vaf*_*j*_:

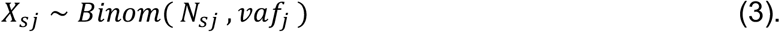

In (2), for a given variant *j*, the expected variant allele frequency,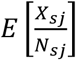, follows a linear relationship the spot deconvolved cell type proportion. The slope Π_*kj*_(*θ*_1*j*_ *– θ*_0*j*_), which is effect size in the regression, and represents the carrier cell type proportion times the difference of allele frequency between real signal and background signal.

To determine the discovery power of real signals under low coverage, we simulated a range of effect sizes (default {0.1,0.3,0.5,0.7,0.9}). For each effect size, we assumed the variant is real and sampled spot coverage *N*_s*j*_ of m data points in the dataset under test. For each of n repetitions (default m=100, n=10), we simulated the observed alternative allele counts using Equation (3). Then a linear regression test is performed on the m sampled data points for n times.

The corresponding power at each effect size is calculated as:

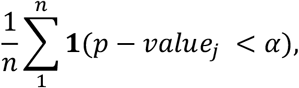

where the default significance threshold α is 0.05.

### Localized Spot-Level Significance Test

While the global cell type test and power analysis help to identify the specific cell types that are co-enriched with each mitochondrial variant, some variants could be too rare and thus have low power. However, when such low frequency variants localize in space, we can leverage their spatial clustering to boost power. We designed a localized spot-level significance test for this purpose.

For each spot s, we first identify its k nearest neighbors set *S*_*s*_ based on spatial Euclidean distance with default k = 100. For each spot s, given variant j and cell type k, we performed a regression of the observed VAFs 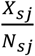 on the deconvoluted cell type proportions *W*_*sk*_, using the neighboring spots *S*_*s*_ and a set of control spots. Control spots are defined as those with sufficient coverage and low proportions of the cell type under consideration (default >=2 coverage and < 0.3 proportion).

As in the global cell type test, spots with significant p-values and positive slope Π_*kj*_(*θ*_1*j*_ *– θ*_0 *j*_) > 0 are identified as significant for variant j and cell type k. Since, multiple spots were tested, we need to control for multiple testing across spots. We allow users to control p-values of the spots via standard Benjamini Hochberg procedure.

In the visualization, we plotted

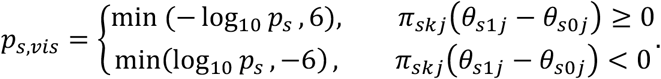

That is, we capped the *−* log_10_ *p*_*s*_ at 6, and assigned p-values with negative slope to be negative values.

### Simulation of Spatial Clones

We simulated three clones on a 30×30 grid of spatial coordinates, with clone sizes drawn from {10,25,50}. Each clone contains five lineage-specific variants, with additional of 15 variants as noise at a VAF of 0.1.

To examine the effect cell type mixtures, four cell types were simulated, with each corresponding to the clone 1-3 and normal cell type. We simulated combinations of lineage *vaf* ∈{0.1,0.25,0.5,0.75,0.9}, coverage *N* ∈{2,5,8,10} and cell type ratio ∈{0.1,0.25,0.5,0.75}.

Given fixed coverage N and VAF, the observed alternative allele count was sampled from:

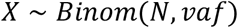

Given spatial coordinates of the cell types, the celltype ratio were simulated from:

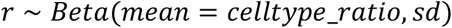

With standard deviation = 0.2.

To measure the performance, we calculated metrics including overall accuracies, class average accuracy, weighted F1 score, Type I error and Type II error.

### Spiking-In of Spatial Clones from Real SUMMIT Data

To represent realistic biological structures, the simulation was also performed based on SUMMIT profiling of the Barrett’s Esophagus sample. Total of three clones were synthesized, with the center spot of each clone manually picked within the Barrett’s Esophagus region of the whole slide. Clone sizes *m* ∈{5, 25, 50,100} and spatial lineage spread *l* ∈{0,1,2} were simulated. For a given center spot *s* with coordinates (*x*_c_, *y*_c_), the set S - the set of spots belonging to the clone, with |*S*| = *m* - were picked as the nearest neighbors of s (*l* = 0), 2^nd^ order nearest neighbors (*l* = 1) and 3^rd^ order neighbors (*l* = 2).

For each variant *j*, define bulk level VAF as p_-_, and define each spot *s* ∈*S* = {1,2, …, *n*}

The observed mitochondrial read depth/coverage matrix *N*_s_ of each spot was directly simulated based on SUMMIT profiling of the Barret Esophagus’ sample:

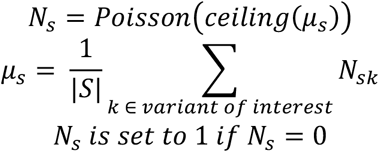

The variant allele frequency *p*_*j*_ was sampled from:

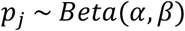

with mean VAF from {0.1,0.25,0.5,0.75,0.9}, and standard deviation = 0.2.

For variant *j* in cell/spot s, the observed mitochondrial variant count matrix *X*_*s j*_ can be synthesized by:

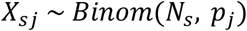

We assumed 1 informative variant per clone and 1 noise variant with background vaf = 0.1 and sd=0.2.

### Spatial Intermixing Index

To determine whether a pair of clones is spatially intermixed or separated, we designed the Spatial Intermixing Index score based on Local Inverse Simpson’s Index (LISI)^25^.

For a pair of variants *i* and *j*, we first filtered for spots that contain sufficient coverage for variant *i* (default coverage *N*_s*i*_ > 0) and with variant *i* detected (*vaf*_s*i*_ > 0). Then for each spot s among these spots, we identified its *k* nearest neighbors 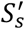 (default k=50).

The intermixing score of spot s represents the number of types of clones detected among its neighboring spots. It is defined as *C*_s,*ij*_ ∈{0,1,2} based on the number of distinct variant types (i or j) detected in neighbors 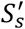.

We also repeated the same process for variant *j*, filtered by coverage and variant allele frequency and get *C*_s,*ji*_.The final LISI score 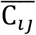 of pair of variants i,j is the average score across all filtered spots:

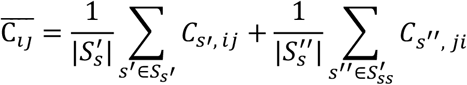

When 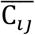 is close to 1, it indicates that the pair of variants are spatially separated, while 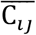 close to 2 suggests the pair of variants are spatially intermixed.

### Spatial Clustering Index

For each variant of interest, we want to determine whether it is spatially clustered. We designed the spatial clustering index based on Moran’s I statistics.

For each variant j, we filtered for spots that contain sufficient coverage for variant *j* (default coverage *N*_s*j*_ > 2) and with variant *j* detected (*vaf*_s*j*_ > 0). For each spot s in this set, we identified the k nearest neighbors 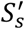 (k=50 by default). The Moran’s I score for variant j is calculated as:

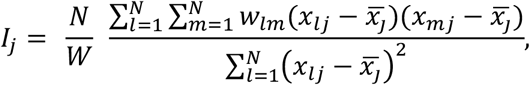

where N is the total number of filtered spots. *x*_*lj*_, *x*_*mj*_ are the variant allele frequencies of spot *l* and spot *m* for variant j, respectively.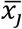 is the average allele frequency of variant j across all the N spots. *w*_*l;*_ represents the spatial weight between spot *l* and spot *m*.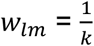, if *m* is the neighbor of spot *l*, otherwise 0. *w*_*ll*_, the weight to self, is set to 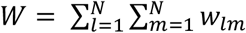, which is the sum of all weights.

The Moran’s I score was calculated using R’s *spdep* package, which also performed Moran’s test. The null hypothesis of the test is that the clone has no spatial autocorrelation (equivalent as randomly scattered in space). P-values from Moran’s test was reported in the Supplementary Table. As a control, we also generated a spatially randomly scattered variant, by resampling variant allele frequency from one variant of interest, while keeping coverage fixed. We calculated its Moran’s I as the control value.

### gDNA extraction and mitochondrial whole exome sequencing

Fresh frozen tissue was equilibrated to -20°C for 20 minutes before sectioning. OCT-embedded tissue was cut at 20 µm thickness in a cryostat. gDNA was extracted from three scrolls per sample using the QIAamp DNA Mini Kit (Qiagen, 51304). For PDAC cell lines Capan-2 and SW1990, gDNA was extracted from cells not injected into the NSG mouse. 50 ng of gDNA per sample was processed with the Twist Mitochondrial Whole Exome Panel (Twist Biosciences, 102038). Indexed libraries were sequenced on an Illumina Nextseq 550 (150 cycles for Read 1/2, 10 cycles per index).

### Whole-exome sequencing data preprocessing, somatic variant calling

FASTQ files were processed according to GATK’s Best Practices. Reads were aligned to the GRch38 reference genome, duplicates were removed, and preprocessed data was analyzed with Mutect2’s mitochondrial model (GATK, v4.2.5.0) for mitochondrial single-nucleotide polymorphism identification. Available germline resource somatic-hg38_af-only-gnomad.hg38.vcf.gz (GATK) served as a matched normal. Variants were filtered using FilterMutectCalls (GATK) and compared to high quality variants detected by maegatk. All variants were cross referenced with known mutations in nuclear mitochondrial DNA and removed if applicable^35^.

### Artifactual analysis in sequencing of the enriched mitochondrial library

We confirmed that detected variants were not artifactual by keeping variants in which 1) the average mtRNA reads per bead for each spatial lineage passed a threshold of >3, 2) the relative position of detected variants were not at the edges of the read, where prior work has reported an increased incidence of sequencing artifacts^50^, and 3) the variant loci was not located within 10 base pairs of the the primer sites on the mitochondrial genome. All spatial variants exceeded a minimum average coverage of 3 reads, as established by Miller et al.l^20^, and were found in a minimum of 100 cells (Fig. S7A). Furthermore, no spatial variant was situated within 20 bases of the read ends, nor within 15 bases of primer binding sites (Fig. S7B-C).

### Use of Artificial Intelligence in manuscript preparation

We enhanced the readability and clarity of this manuscript with assistance from Claude (Anthropic, San Francisco, CA), a large language model artificial intelligence system. Claude was used to check grammar and spelling, improve sentence structure, and overall readability of the text while preserving the original scientific content and conclusions. Suggested revisions were reviewed and improved by the authors before incorporation into the manuscript. The use of AI assistance was limited to editorial improvements and did not involve generation of research data, statistical analyses, or scientific conclusions. Data collection, analysis, and interpretation was the responsibility of the authors, independent of artificial intelligence.

## Author contributions

Conceptualization – S.A.B., R.A.G., and S.M.S.; Methodology – S.A.B., R.A.G., J.R., N.R.Z., and S.M.S.; Investigation – S.A.B. and J.R.; Software – S.A.B. and J.R.; Formal Analysis – S.A.B., J.R., and N.R.Z.; Data Curation – S.A.B.; Mouse colony maintenance and xenograft generation – J.C.M., F.M., and B.Z.S.; Writing – Original Draft, S.A.B., J.R., N.R.Z. and S.M.S.; Writing – Review and Editing, S.A.B., J.R., N.R.Z. and S.M.S.; Visualization – S.A.B., J.R., N.R.Z. and S.M.S.; Resources – H.G, D.D, M.D., G.W.F., B.W.K., E.E.F., A.B.M., and A.S.S.; Funding Acquisition – S.M.S.; Supervision – N.R.Z. and S.M.S.

## Acknowledgements

We thank the members of the Shaffer and Zhang groups for input on the methodology and figures of the manuscript. This work was supported by National Science Foundation grant DMS-2245575 and National Institutes of Health (NIH) grant 1R56AG081351 to J.R. and N.R.Z. Additional support was provided by NIH grant 5R01GM14967 to S.M.S. and N.R.Z., NIH grant 1R01DK135729 to S.M.S., N.R.Z., and A.B.M., and NIH grants R01CA252225 and R01CA276512 to B.Z.S. The work was also supported by the University of Pennsylvania Basser Center Internal Grant to S.M.S. and B.K. S.M.S. is the Bakewell Foundation Innovator of the Damon Runyon Cancer Research Foundation (DRR-81-24) and is further supported by American Cancer Society Research Scholar Grant RSG-23-1152597-01-CDP.

## Conflicts of interest

All authors have no conflicts of interest to declare.

## Notes

### Competing Interest Statement

The authors have declared no competing interest.

