## Supplemental figures for "Mitochondrial clone tracing within spatially intact human tissues"

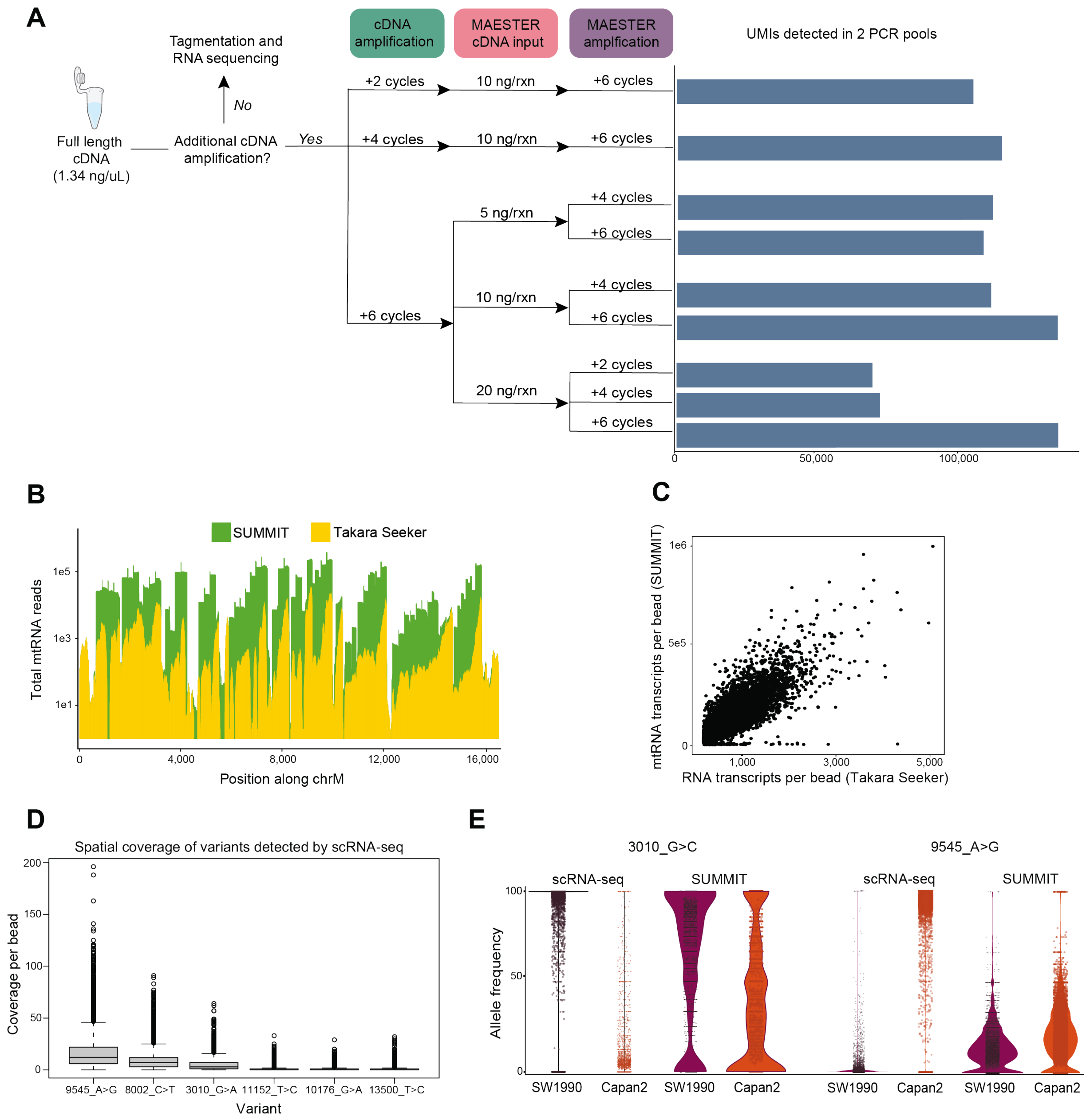


**Figure S1: Optimization and validation of the SUMMIT workflow.** (A) Systematic optimization of SUMMIT protocol parameters. Decision tree showing optimization of cDNA amplification cycles, input amounts, and mitochondrial-specific PCR cycles to maximize mitochondrial transcript recovery. Bar graph shows UMI detection across parameter combinations. (B) Comparison of mitochondrial transcript coverage between Takara Seeker libraries (yellow) and SUMMIT-enriched libraries (green) across the mitochondrial genome, demonstrating 25- to 100-fold enrichment. (C) Scatter plot showing correlation between mitochondrial transcript coverage in SUMMIT and Takara Seeker libraries, confirming efficient enrichment without substantial bias. (D) Coverage comparison for cell type specific variants detected by single-cell RNA sequencing, showing variable detection sensitivity for different variants in SUMMIT data. (E) Violin plots comparing allele frequencies for variants 3010_G>C and 9545_A>G between cell lines in both single-cell and SUMMIT datasets.

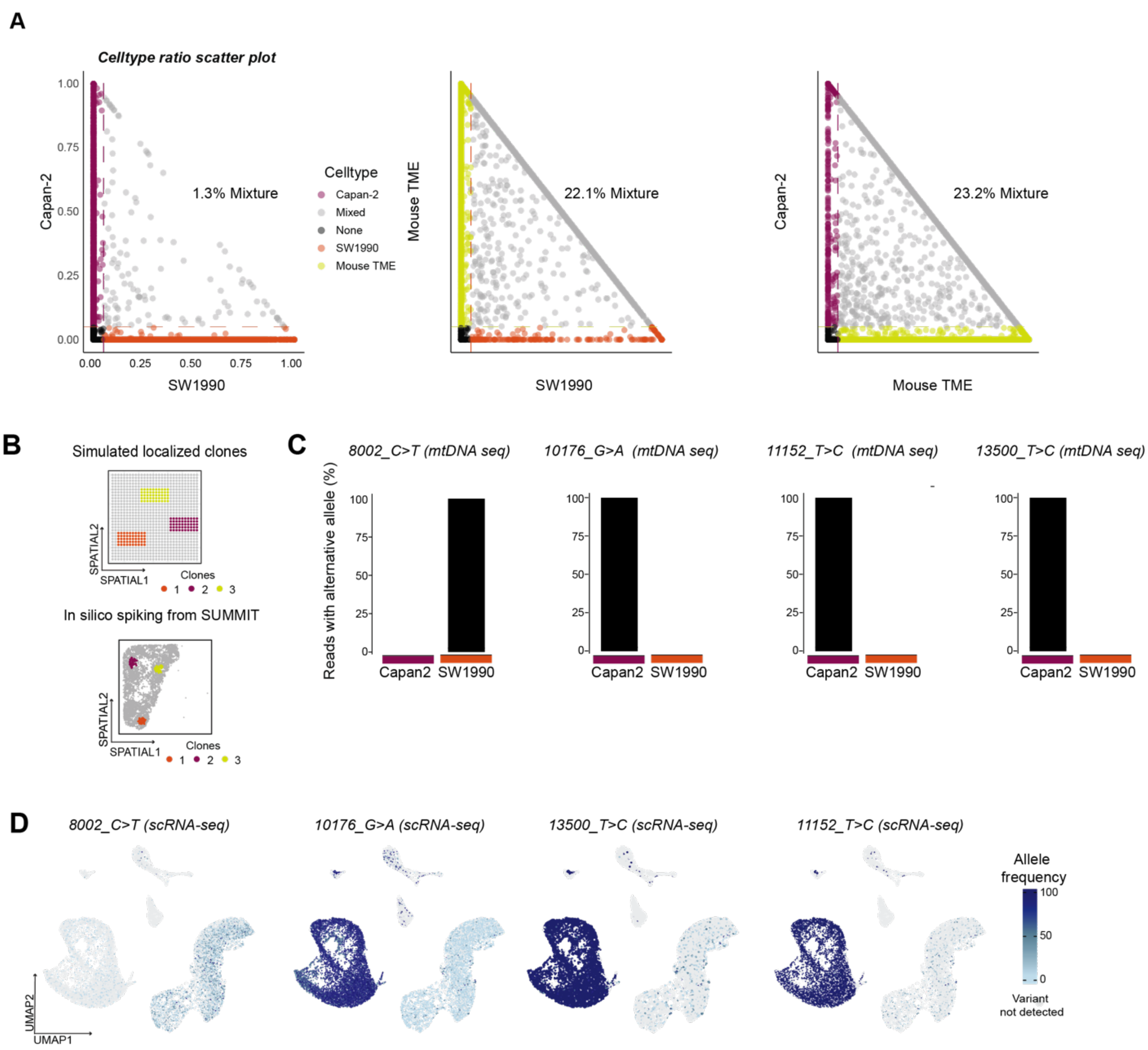


**Figure 2S: Validation of cell type mixing and computational pipeline performance in xenograft experiment.**

(A) Scatterplots of cell type ratio showing the distribution of cell type proportions for Capan-2, SW1990, and murine cells within spatial spots for the xenograft experiment. Each plot displays the proportion of one cell type versus another, with colors indicating the dominant cell type assignment. Percentages indicate the fraction of spots containing mixtures of the specified cell types. (B) Simulation framework designs used to evaluate SUMMIT's computational pipeline. Top: Simulated localized clones on a 30×30 grid with 3 clones each carrying distinct variants. Bottom: In silico spiking approach using real SUMMIT data structure to test detection under varying biological parameters. (C) mtDNA sequencing validation of cell line-specific variants detected in the xenograft experiment. Bar plots show the percentage of reads with alternative alleles for variants 8002_C>T, 10176_G>A, 11152_T>C, and 13500_T>C in Capan-2 (purple) and SW1990 (orange) cell lines, confirming cell line specificity of detected variants. (D) Single-cell RNA sequencing validation showing cell type specificity of iants 8002_C>T, 10176_G>A, 11152_T>C, and 13500_T>C in UMAP space.

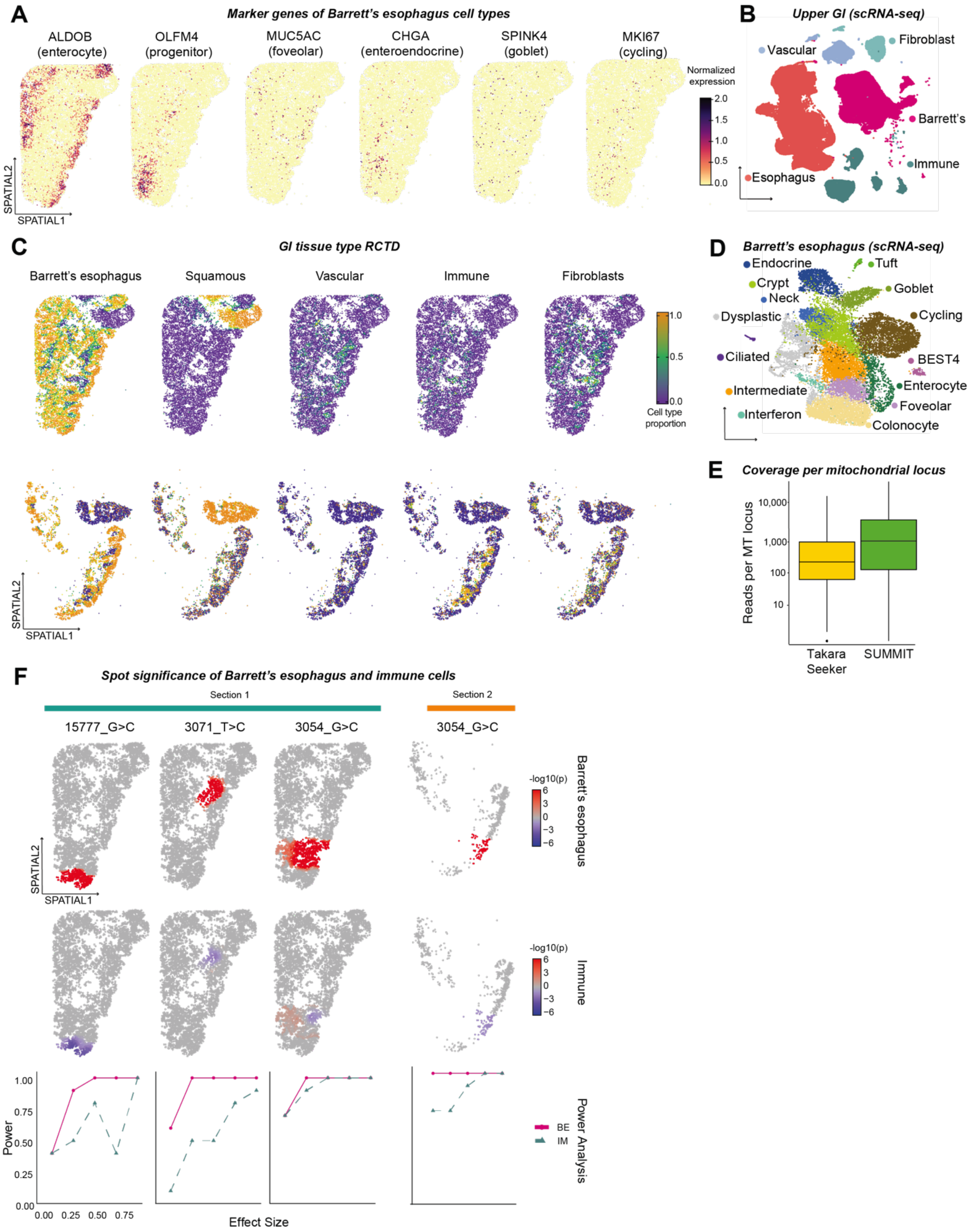


**Figure S3: Detailed characterization of Barrett's esophagus spatial architecture and mitochondrial clone detection.**

(A) Spatial gene expression patterns of Barrett's esophagus cell type markers including ALDOB (enterocyte), OLFM4 (progenitor), MUC5AC (foveolar), CHGA (enteroendocrine), SPINK4 (goblet), and MKI67 (cycling). (B) Single-cell RNA sequencing reference atlas[27] of upper gastrointestinal tissue showing distinct clustering of fibroblast, vascular, Barrett's esophagus, squamous, and immune populations. (C) Robust cell type deconvolution (RCTD) maps showing spatial distribution of Barrett's esophagus, squamous, vascular, immune, and fibroblast populations with corresponding cell type proportion scales. (D) Single cell RNA-sequencing atlas[27] used for Barrett's esophagus-specific cell type annotation. (E) Coverage comparison between Takara Seeker and SUMMIT libraries showing ~10-fold increase in mitochondrial transcript detection. (F) Spot-level significance testing visualization for spatially-confined variants across tissue sections, showing spatial patterns of significance for Barrett's esophagus associations. Corresponding power analysis curves demonstrate adequate statistical power for confident negative results.

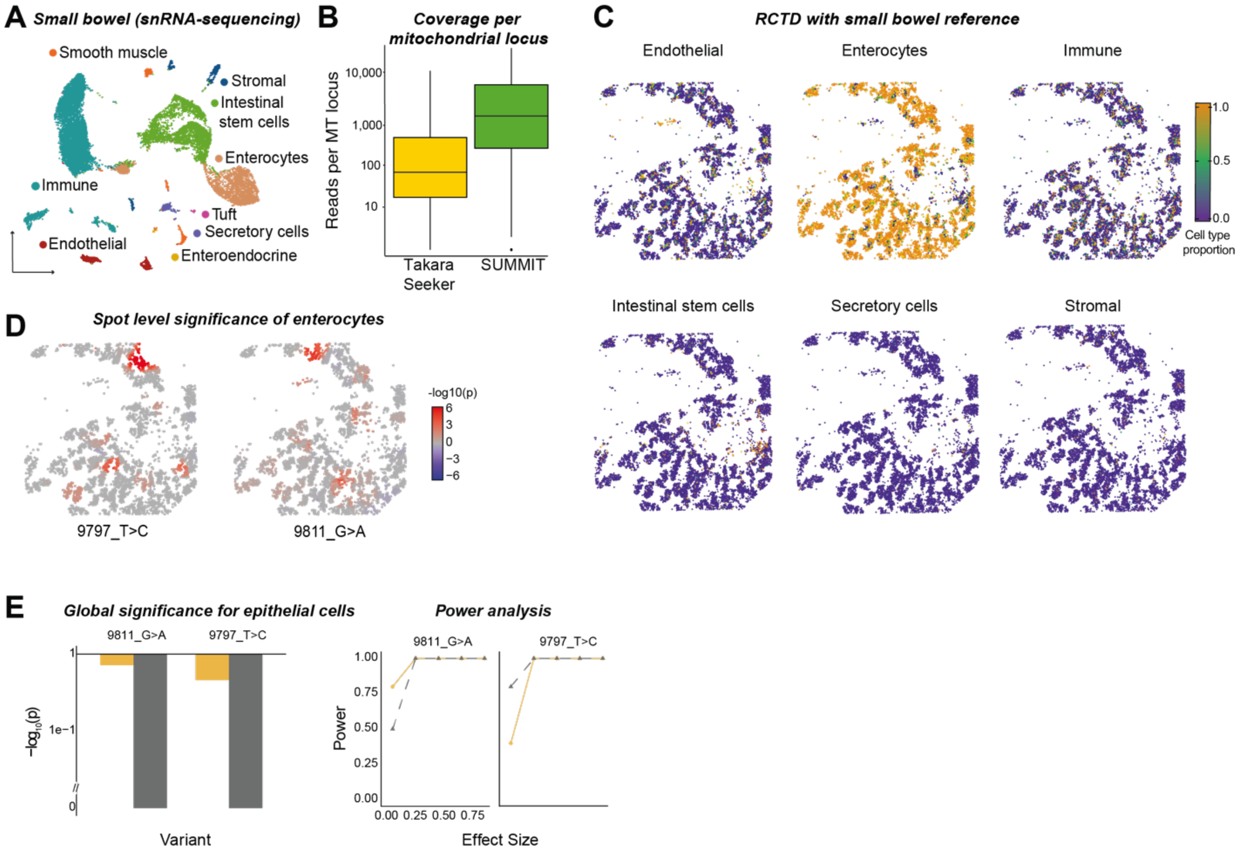


**Figure S4: Spatial characterization and power analysis of independent clones in normal small bowel.** (A) Single-nucleus RNA sequencing reference dataset contains distinct intestinal cell types including intestinal stem cells, enterocytes, tuft cells, secretory cells, and enteroendocrine cells. (B) Coverage comparison between Takara Seeker and SUMMIT libraries demonstrating significant improvement in mitochondrial transcript detection. (C) RCTD maps showing spatial distribution of endothelial, enterocytes, immune, intestinal stem cells, secretory cells, and stromal populations across the tissue section. (D) Spot-level significance testing visualization for variants 9797_T>C and 9811_G>A, showing spatial patterning of significance for enterocytes. (E) Global significance testing and power analysis for epithelial cells, demonstrating lack of significant association of both variants with intestinal epithelium and immune cells, while reporting adequate statistical power for detecting variants in immune cells if present.

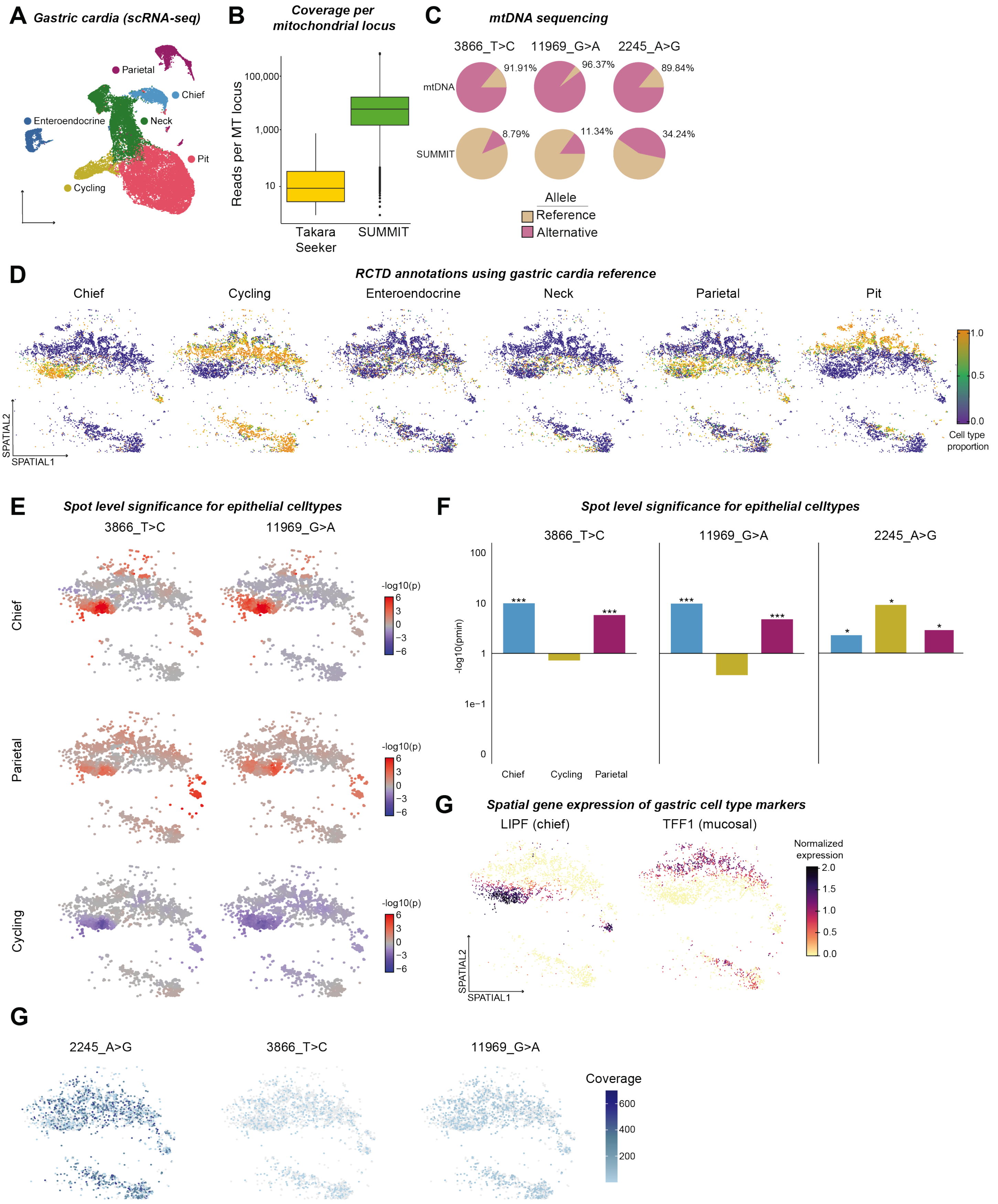


**Figure S5: Detailed spatial analysis of normal gastric epithelium and subclonal relationships.**

(A) Single-cell RNA sequencing reference atlas[27] of gastric cardia showing distinct gastric cell types including parietal, chief, neck, and pit cells. (B) Coverage comparison between Takara Seeker and SUMMIT libraries showing improvement in mitochondrial transcript detection. (C) mtDNA sequencing validating all three variants detected by SUMMIT. (D) RCTD maps showing spatial distribution of chief, cycling, enteroendocrine, neck, parietal, and pit cell populations across the gastric tissue section. (E) Spot-level significance testing visualization for epithelial cell subtypes showing spatial patterns of significance for chief and parietal cell associations with variants 3866_T>C and 11969_G>A. (F) Global significance testing across gastric epithelial subtypes confirming significant associations with adequate statistical power. (G) Spatial gene expression maps for gastric cell type markers *LIPF* (chief) and *TFF1* (mucosal), showing correspondence between marker expression and clone locations.

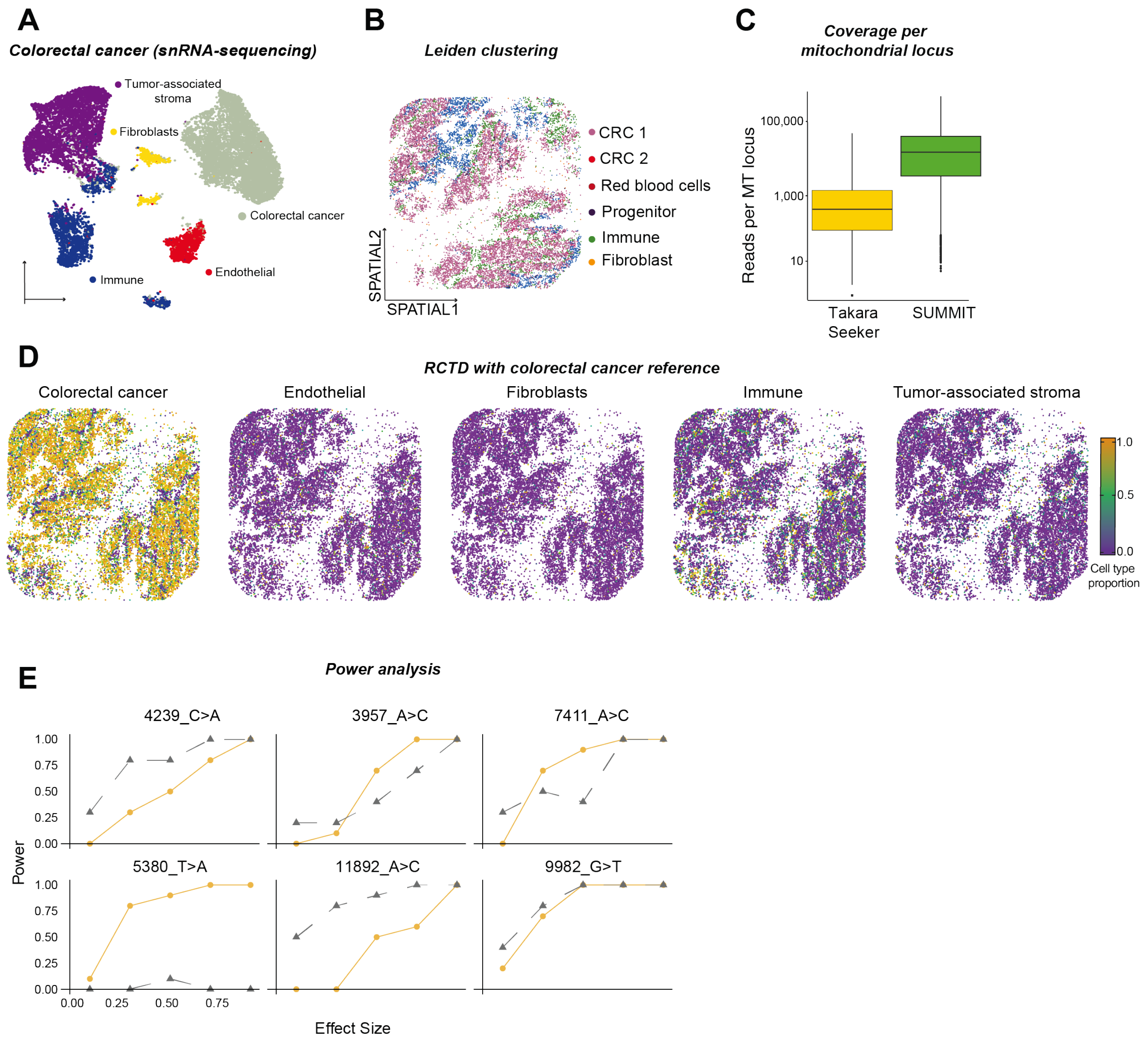


**Figure S6: Detailed characterization of colorectal cancer spatial architecture and intermixed clonal populations.** (A) Single-nucleus RNA sequencing reference atlas showing clustering of colorectal cancer epithelial populations, immune cells, and endothelial populations. (B) Leiden clustering of spatial transcriptomics data showing distinct colorectal cancer clusters (CRC 1 and CRC 2) along with red blood cells, progenitor, immune, and fibroblast populations. (C) Coverage comparison between Takara Seeker and SUMMIT libraries demonstrating significant improvement in mitochondrial transcript detection. (D) RCTD maps showing distribution of colorectal cancer, endothelial, fibroblast, immune, and tumor-associated stroma populations across the tissue section. (E) Power analysis for all six variants showing adequate statistical power for detection across different effect sizes, confirming reliability of significance testing results.

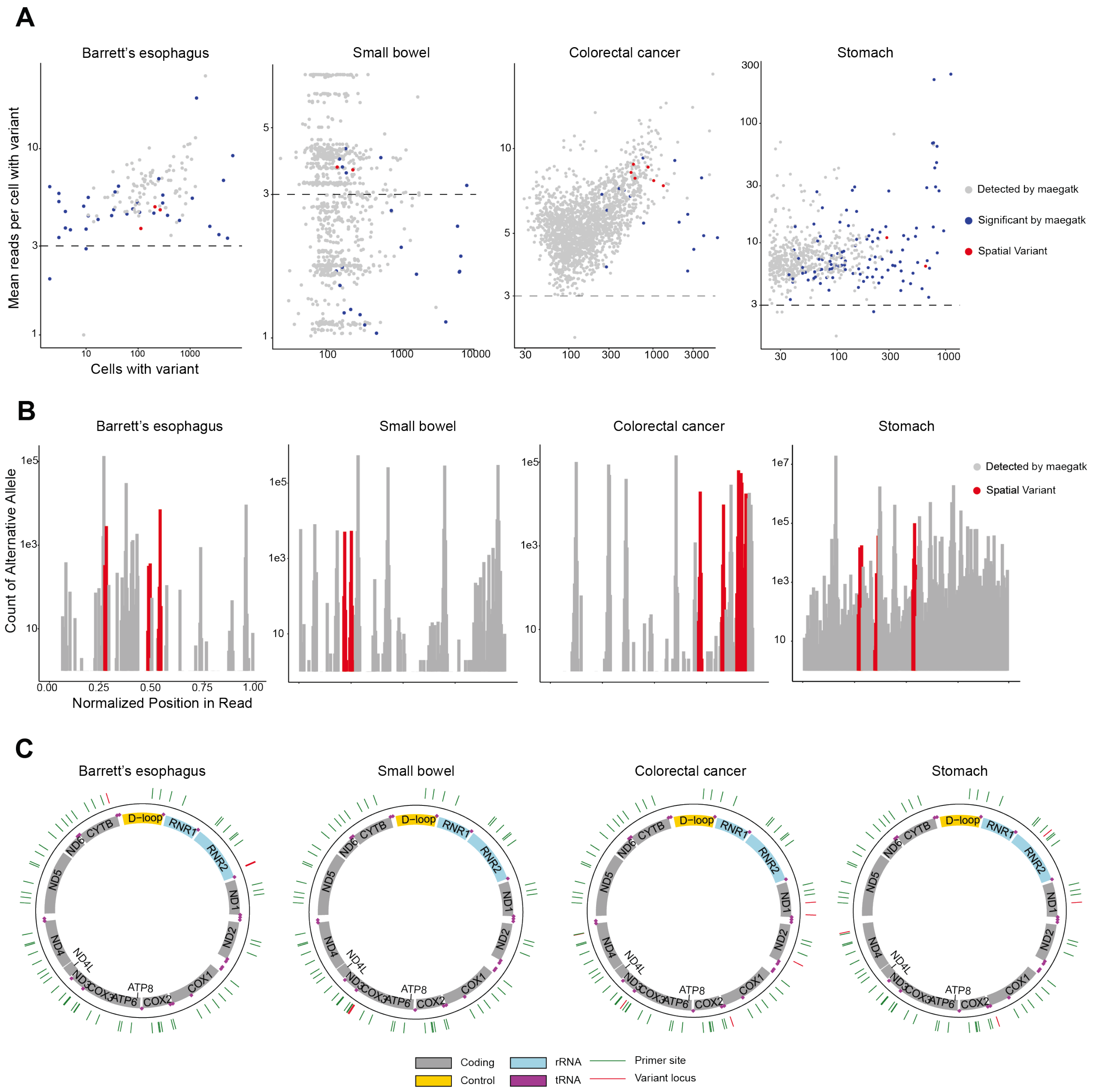


**Figure S7: Technical validation of mitochondrial variant detection and sequencing artifact analysis.** (A) Cross-tissue comparison of variant detection showing mean reads per cell with variant size. Variants detected by SUMMIT are colored in blue, with spatial variants shown in red. Dashed lines indicate detection thresholds. (B) Normalized position analysis in sequencing reads showing distribution of detected variants across read length, confirming variants are not preferentially detected at read ends where sequencing artifacts are common. (C) Mitochondrial genome maps showing locations of detected variants (red bars) relative to mitochondrial genes, primer binding sites, and control regions. Analysis confirms variants are not located near primer sites.
